# Brain-wide reconfiguration of burst firing by psilocybin reveals 5-HT2A-dependent circuit dynamics

**DOI:** 10.64898/2026.08.14.744865

**Authors:** D. Momi, Y. Nahas, D. Wyrick, L. C. Marks, L. D. Claar, R. De Filippo, P. Seyfourian, D. A. Pizzagalli, M. Buice, T. Ott, C. Koch, I. Rembado

## Abstract

Psilocybin produces rapid and lasting therapeutic effects, yet how 5-HT2A receptor activation reshapes brain-wide circuit dynamics during acute drug administration remains poorly understood. Using simultaneous multi-region Neuropixels recordings of 46,360 single units from 35 mice, together with scalp electroencephalography (EEG), pupillometry, and locomotion monitoring, we provide a brain-wide, single-unit and field-potential characterization of psilocybin’s acute effects, with pharmacological dissection using the 5-HT2A antagonist ketanserin. Psilocybin selectively reconfigured burst coding, rather than mean firing rate, across cortical, thalamic, and hippocampal circuits: burst firing decreased in hippocampal CA1–CA3 and was bidirectionally modulated in the thalamus, with the reticular nucleus bursting more and first-order geniculate nuclei bursting less. Critically, most of these burst effects were abolished by ketanserin, consistent with at least partial 5-HT2A receptor dependence. These data suggest that the psychedelic state is not simply a matter of how much neurons fire, but of how they fire, pointing to a region-specific, 5-HT2A-associated reconfiguration of burst coding that may underlie the acute phenomenology of the psilocybin experience.

## INTRODUCTION

A single dose of psilocybin in humans can produce lasting relief from depression, anxiety, and substance use disorders (Vollenweider and Preller 2020; Nutt, Erritzoe, and Carhart-Harris 2020; Raison et al. 2023), an effect not typically observed with conventional psychiatric medications, whose benefits require continuous dosing, have sustained side effects and disappear once treatment stops. This durability is thought to rest on the induction of structural plasticity: through partial agonism at serotonin 2A receptors (5-HT2ARs), densely expressed on the apical dendrites of cortical layer 5 pyramidal neurons in mice (Vollenweider et al. 1998; Lopez-Gimenez and Gonzalez-Maeso 2018), psilocybin drives rapid dendritic spine growth in prefrontal cortex (Shao et al. 2025; Shao et al. 2021), activity-dependent rewiring of cortical networks (Jiang et al. 2026), hippocampal synaptic plasticity (Hesselgrave et al. 2021; Raval et al. 2021), and shifts in functional connectivity (Daws et al. 2022) that outlast the acute effect of the drug itself. Yet plasticity is a downstream consequence: what psilocybin does to neural activity during the hours it is pharmacologically active, and how that acute reconfiguration gives rise to the lasting therapeutic effects, remains largely unknown.

As little is known about the action of psilocybin at the cellular levels in humans, pre-clinical models are critical. So far, most accounts of psychedelic action in mice rest on reports of changes in mean firing rate and oscillatory power (Hidalgo Jimenez, Kristjan Kaup, and Aru 2026; Purple et al. 2025; Golden and Chadderton 2022; Skyberg et al. 2025; Breant et al. 2026); yet rate is only one axis along which neurons carry information. Bursts - rapid successions of spikes at short interspike intervals - drive postsynaptic targets far more reliably than isolated spikes, trigger supralinear dendritic summation and calcium influx, and are specifically required for burst-dependent long-term potentiation (Lisman 1997; Magee and Johnston 1997; Harris et al. 2001; Thomas et al. 1998; Pike et al. 1999; Koch 1999). Thus, they have been proposed to constitute a coding channel distinct from single-spike firing, signaling the coincidence of bottom-up and top-down input (Larkum 2013; Lisman 1997; Bair et al. 1994). Because this coincidence-detection role ties burst generation to the balance between bottom-up sensory drive and top-down contextual input, bursts may be especially sensitive to the acute reorganization of cortical processing that psychedelics are thought to produce (Carhart-Harris and Friston 2019; Vollenweider and Geyer 2001; Doss et al. 2022; Corlett et al. 2019; Kwan et al. 2022). Whichever direction the imbalance takes, bursts should be sensitive to it. If psilocybin acts acutely by rebalancing these input streams rather than by uniformly scaling excitability, the relevant signature should lie not in how much neurons fire, but in how they fire; i.e., their burst dynamics rather than their overall firing rate.

Testing these predictions requires sampling neural activity broadly across the brain. Human neuroimaging localizes psilocybin’s action in the cortico-striato-thalamo-cortical (CSTC) circuit, whose dysregulation, through thalamocortical dysrhythmia at its thalamic gate, underlies several psychiatric disorders (Alkire, Hudetz, and Tononi 2008; Crick 1984; Geyer and Vollenweider 2008; Ward 2011; Ferrarelli and Tononi 2011; Llinas et al. 1999; Onofrj et al. 2023). Psilocybin alters CSTC and default mode network dynamics, but studies disagree on the direction of this effect, reporting both increased and decreased bottom-up processing (Carhart-Harris et al. 2012; Preller et al. 2019; Gaddis et al. 2022; Pines et al. 2026). This disagreement is likely a resolution problem rather than a true absence of a canonical response, since imaging treats activity as stationary and aggregates signals across too many neurons to resolve either directionality or cellular origin (Pines et al. 2026; Logothetis 2008). Yet the cellular recordings capable of that resolution have been confined almost entirely to single regions, predominantly prefrontal cortex in rodents (Hidalgo Jimenez, Kristjan Kaup, and Aru 2026; Vollenweider and Preller 2020). No study has recorded single neurons simultaneously across these distributed nodes during the acute phase of psilocybin in awake, behaving animals.

To isolate which of psilocybin’s acute effects on neural activity depend on 5-HT2A receptor activation, we used ketanserin, a 5-HT2A/2C antagonist that abolishes psilocybin’s perceptual effects in humans (Vollenweider et al. 1998; Kometer et al. 2013) and the head-twitch response in rodents (Halberstadt 2015; Hesselgrave et al. 2021). Here, we combined this pharmacological dissection with brain-wide recording: across 47 sessions in awake, head-fixed mice, we measured >46,000 single units and local field potentials simultaneously across cortical and subcortical CSTC nodes and the hippocampus during the acute effects of psilocybin, alongside scalp EEG, pupillometry, and locomotion. We found that psilocybin selectively reconfigured burst coding, rather than mean firing rate, across the CSTC circuit and hippocampus in a region-specific pattern consistent with a rebalancing of top-down and bottom-up signaling. This reconfiguration was abolished by ketanserin, indicating at least partial 5-HT2A receptor dependence.

## RESULTS

### Multiscale electrophysiological and behavioral recordings during acute psilocybin administration in head-fixed mice

We performed head-fixed recordings in mice while simultaneously monitoring locomotion, pupil dynamics, electroencephalography (EEG), local field potentials (LFPs), and single-unit activity. During each session, mice were head-fixed on a rotating wheel, free to rest or run, for approximately 2 hours (Figure 1a, S1a-d). Wheel angular velocity was recorded to track locomotion speed, and an infra-red video of the right eye, reflected by a dichroic mirror, was acquired to track pupil size (Figure 1a, S1a-d). EEG signals were recorded using a 30-channel array chronically implanted directly on the skull, below the scalp (Figure 1b, top), and extracellular signals were acquired using up to four Neuropixels 1.0 probes (Jun et al. 2017) spanning cortical and subcortical regions (Figure 1b, bottom; Figure 1e). During each experimental session, mice received two intraperitoneal (i.p.) injections spaced 10–12 min apart while head-fixed (Figure 1d). In saline control sessions, both injections were saline (0.01 mL/g at 0.9%). In psilocybin sessions, the first injection was saline and the second was psilocybin (1 mg/kg). In ketanserin+psilocybin sessions (referred to as Ket+Psi hereafter), the first injection was ketanserin (1 mg/kg), and the second was psilocybin (1 mg/kg). The period prior to any pharmaceutical intervention (ranging from 15 to 45 min depending on session protocol) was used as baseline for behavioral measurements; for unit analyses, the baseline was restricted to the last 10 min before the first injection to ensure consistency across animals with different pre-injection periods. The 60 min period following the second injection was used to compare experimental conditions; as it falls within the acute behavioral window of psilocybin at this dose, which produces head-twitch responses peaking at around 8 min and lasting up to around 2 h post-injection (Shao et al. 2021). Across 22 psilocybin, 17 saline, and 8 Ket+Psi sessions in 35 animals, we recorded 46,360 single units, comprising 38,429 regular-spiking (RS) and 7,931 fast-spiking (FS) units (see Methods for classification details) distributed across motor, somatosensory, visual, and prefrontal cortical areas, as well as subcortical regions including multiple thalamic nuclei, striatum, and hippocampus (for a breakdown of units count per area see Supplementary Table 1; Figure 1c). Psilocybin had no significant effect on locomotion speed across any condition (Figure S1e–f; Wilcoxon signed-rank tests, p > 0.05), consistent with previous studies (Fadahunsi et al. 2022; Lu et al. 2025; Purple et al. 2025). In contrast, when controlling for locomotion speed and time in the experiment with a linear mixed-effects model in paired sessions, psilocybin induced a significant increase in pupil size relative to baseline (Figure S1g–j), confirming the behavioral effects we observed on a separate and independent dataset (De Filippo et al. Under submission). Crucially, pupil dilation was also present in Ket+Psi sessions, indicating that this effect was not abolished by ketanserin pretreatment (Figure S1k–l).

**Figure 1.**
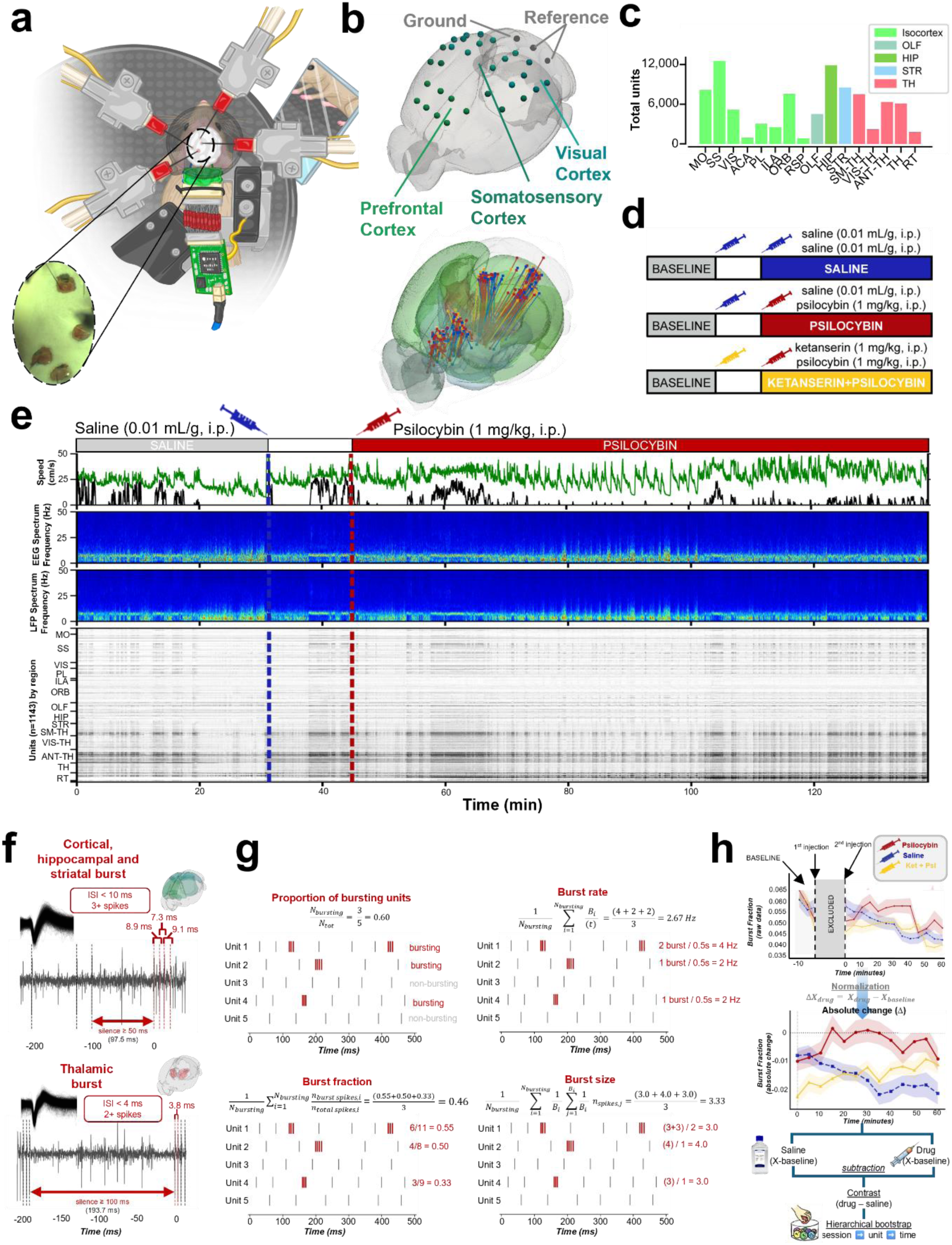
Multiscale electrophysiological and behavioral experimental setup in head-fixed mice. a, Graphic of the experimental setup showing the head-fixed mouse on a rotating wheel used to monitor locomotion, a dichroic mirror used to monitor the eye for pupillometry, the electroencephalogram (EEG) electrodes on top of the skull (beneath the white cement), and four Neuropixels 1.0 probes inserted into craniotomies in the left hemisphere. b, Top: Schematic of the 30-channel EEG array (green circles) implanted on the skull surface over major brain areas, including prefrontal, somatosensory, and visual regions. Also shown are three skull screws over the cerebellum (gray circles) serving as the reference and ground for both EEG and Neuropixels signals. Bottom: Neuropixels probe traces across all subjects included in this dataset (47 sessions across 35 mice), targeting cortical and subcortical areas shown in a reconstructed common coordinate framework. Probe traces are colored by experimental condition: saline (blue), psilocybin (red), and ketanserin+psilocybin (yellow), as in d. Schematics were created using BrainRender (Claudi et al., 2021). c, Total number of units isolated from each brain region across all subjects in this dataset (47 sessions across 35 mice). d, Experimental timeline for saline control (top), psilocybin (middle), and ketanserin+psilocybin control (bottom) experiments. In each experiment, the subject received two intraperitoneal (i.p.) injections approximately 10 min apart, as indicated by the syringe icons; the contents of each injection are listed to the right. e, Experimental timeline, behavioral data, and multiscale electrophysiological recordings from a representative psilocybin experiment. The first injection of saline (0.01 mL/g, i.p.; vertical dashed blue line) was administered immediately following a 15-45 min baseline period, and the second injection of psilocybin (1 mg/kg, i.p.; vertical dashed red line) was given approximately 10 min later; recording continued for approximately 1-1.5 hours after psilocybin administration. Traces shown from top to bottom: locomotion speed (black) and pupil radius (green); average power spectrogram across all EEG channels; average power spectrogram across all cortical LFP channels from the Neuropixels probes; raster plot of all units across major brain regions from this experiment. Region abbreviations used in c and e — MO: motor areas; SS: somatosensory areas; VIS: visual areas; ACA: anterior cingulate area; PL: prelimbic area; ILA: infralimbic area; ORB: orbital area; RSP: retrosplenial area; OLF: olfactory areas; HIP: hippocampal region; STR: striatum; SM-TH: somatomotor thalamic nuclei; VIS-TH: visual thalamic nuclei; ANT-TH: anterior thalamic nuclei; TH: other thalamic nuclei; RT: reticular nucleus of the thalamus; FT/CC: fiber tracts and corpus callosum. f, Top: Example cortical and hippocampal burst. Overlaid spike waveforms are extracted from 20 min of recording (black) with the corresponding AP raw signal shown below. In these areas a burst is defined as ≥ 3 spikes with inter-spike intervals (ISI) < 10 ms (example ISIs: 7.3, 8.9, 9.1 ms), preceded by a silence of ≥ 50 ms (97.5 ms shown). Bottom: Example of thalamic burst. A burst is defined as ≥ 2 spikes with ISI < 4 ms (example ISI: 3.8 ms), preceded by a silence of ≥ 100 ms (193.7 ms shown). Brain schematics indicate the recording regions for each burst type. g, Four burst metrics illustrated on an example population of 5 units from orbital area layer 5 recorded over 500 ms. Red tick marks indicate spikes belonging to bursts; grey ticks indicate non-burst spikes. Proportion of bursting units: fraction of units classified as bursting (≥ 1 burst detected) out of all recorded units (example: 3/5 = 0.60). Burst rate: mean number of bursts per second across bursting units (example: 2.67 Hz). Burst fraction: fraction of spikes occurring within bursts over the total number of spikes, averaged across bursting units (example: 0.46). Burst size: mean number of spikes per burst, averaged across bursts and bursting units (example: 3.33 spikes/burst). h, Normalization and statistical procedure. Top: Example raw burst fraction time course from RS units in layer 5 prefrontal cortex for psilocybin (red), saline (blue), and Ket + Psi (yellow) groups, aligned to the second injection (time 0 s, the shaded gray area indicates the excluded time between the two injections). Color shading indicates standard error of the mean (SEM). Middle: Absolute change (Δ) is computed per unit as the difference between the post-injection value and the unit’s own pre-injection baseline mean (ΔX = X_drug − X_baseline) for each time bin. Bottom: drug–saline contrast is computed by subtracting the saline condition’s absolute change from the drug condition’s absolute change. Statistical significance is assessed using hierarchical bootstrap resampling sessions, then units within sessions, independently at each time bin, generating a null distribution of the contrast under the hypothesis of no drug effect.

To characterize how psilocybin modulated burst firing, we applied region-specific burst definitions, distinguishing thalamic from cortical, hippocampal and striatal bursting (Figure 1f, see Methods for details). We then defined four complementary metrics across all brain regions to quantify burst dynamics: the proportion of units classified as bursting (i.e. firing at least one burst within the analyzed time window), and for each bursting unit, burst rate, burst fraction (fraction of spikes occurring within bursts), and mean burst size (Figure 1g). To quantify drug effects, each metric was expressed as the absolute change from the pre-injection baseline (i.e. prior to the first injection), and a drug–saline contrast was computed and assessed for significance using a hierarchical bootstrap resampling sessions, then units within sessions, independently at each time bin (Figure 1h, see Methods for details).

### Psilocybin bidirectionally modulates burst firing across brain regions partially mediated by 5-HT2A receptor mechanisms

Burst firing was strongly modulated by behavioral state: many neurons that fired in burst mode during quiet rest transitioned to tonic firing during locomotion and arousal (Marlinski and Beloozerova 2014). Consistent with this, a supplementary analysis confirmed that across units, burst fraction was substantially reduced during locomotion in most regions examined (Figure S2). We therefore restricted the burst analysis to periods of rest (speed < 1 cm/s; see Methods), which substantially reduced the number of units contributing to each time bin. The scale of the dataset nonetheless ensured that unit numbers remained large throughout the analyzed time window (Supplementary Table 2).

Illustrating the drug effect at the single-unit level, raster plots from an example psilocybin session showed a marked increase in burst firing in reticular thalamus (RT) units (Figure 2a), with example inter-burst interval (IBI) histograms of representative units showing a corresponding shift toward shorter intervals after psilocybin (2805 ms → 974 ms), a smaller shift under Ket+Psi (334 ms → 287 ms), and minimal change under saline (534 ms → 514 ms) (Figure 2b).

**Figure 2.**
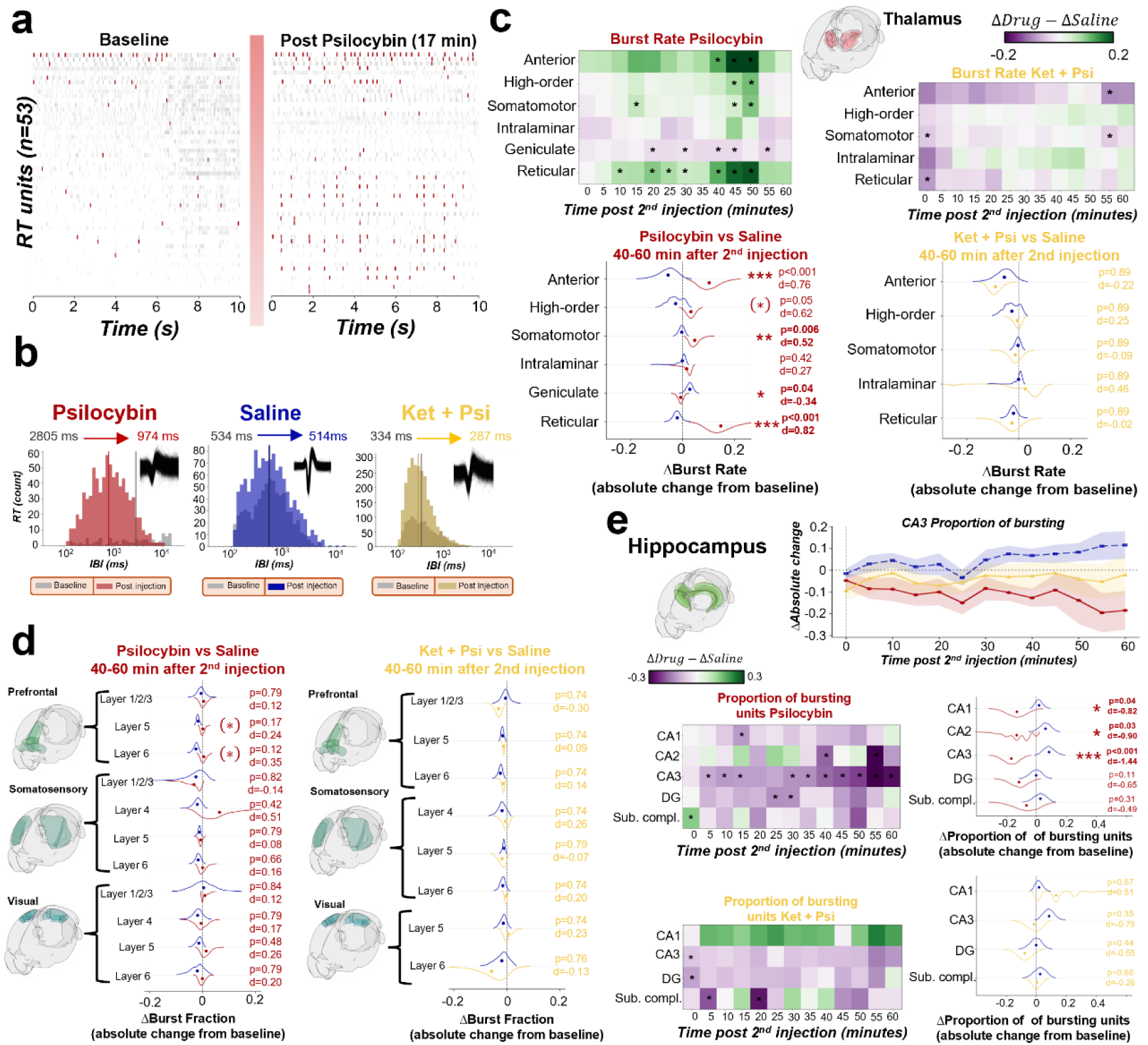
Psilocybin bidirectionally modulates thalamic burst firing and decreases hippocampal bursting, partially mediated by 5-HT2A receptors. a, Raster plot of n=53 reticular nucleus of the thalamus (RT) units from an example psilocybin session, showing 10-second windows during baseline (left) and after psilocybin injection (right) at 17 min after drug administration. Red ticks indicate identified bursts; grey ticks indicate single spikes. b, Inter-burst interval (IBI) histograms for some RT units from example sessions, comparing baseline prior to the first injection (grey) and post-injection distributions (red, psilocybin; blue, saline; yellow, ket+psi). Vertical lines indicate the median of the baseline (grey) and post-injection (colored) distributions. Insets show overlaid spike waveforms of the selected unit from 20 min raw signal. Psilocybin shifts the IBI distribution toward shorter intervals (median: 2805 ms → 974 ms), consistent with increased burst firing. c, Burst rate for thalamic subregions as defined in the methods. Top: Heatmaps showing the drug–saline contrast in burst rate (ΔDrug − ΔSaline) across time post second injection (0–60 min) for psilocybin (left) and Ket + Psi (right). Each row is a thalamic subregion. Asterisks indicate time bins where the contrast significantly differs from zero (hierarchical bootstrap, p < 0.05, uncorrected). Color scale: green, increase; violet, decrease; range ±0.2. Bottom: Half-violin plots showing the distribution of burst rate averaged between 40 and 60 min after second injection for each subregion. Dots indicate the mean; p-values and Cohen’s d are shown after FDR correction. Solid asterisks denote significance after FDR correction (*p < 0.05, **p < 0.01, ***p < 0.001); asterisks in parentheses denote nominal significance (uncorrected p < 0.05) that did not survive FDR correction. Psilocybin significantly increases burst rate in the reticular nucleus (RT; p<0.001, d=0.82), the anterior thalamic nuclei (p<0.001, d=0.76), and the first-order somatomotor nuclei (p=0.006, d=0.52), while significantly decreasing it in the first-order sensory geniculate nuclei (p=0.04, d=−0.34). Psilocybin also induces a trend of increase in higher-order thalamic nuclei that does not survive FDR correction (p=0.05, d=0.62). Ket + Psi produces no significant effects in any thalamic subregion. d, Burst fraction for isocortex subregions by layer. Half-violin plots showing the distribution of burst fraction averaged between 40 and 60 min after second injection for prefrontal (top), somatosensory (middle), and visual (bottom) cortex, stratified by layer. Dots indicate the mean; p-values and Cohen’s d are shown after FDR correction. Left column: Psi vs. saline. Right column: Ket + Psi vs. saline. Burst fraction in deep prefrontal layers decreases from baseline under saline, while psilocybin maintains burst fraction near baseline levels, yielding a drug–saline contrast in layer 5 and layer 6 that does not survive FDR correction, as shown in parentheses. A similar trend is visible in burst rate (Figure S4). No significant drug–saline contrast is observed in somatosensory or visual cortex, or with Ket + Psi in any region or layer. e, Proportion of bursting units in hippocampal areas. Top: time course of the absolute change in proportion of bursting units in CA3 for psilocybin (red), saline (blue), and Ket + Psi (yellow). Shading indicates standard error of the mean (SEM). Bottom left: heatmaps showing the drug–saline contrast in proportion of bursting units across time post second injection for psilocybin (top) and Ket + Psi (bottom) in hippocampal subregions (CA1, CA2, CA3, DG, Subicular complex). Color scale ±0.3. Asterisks indicate significant time bins. Bottom right: Half-violin plots showing the distribution of proportion of bursting units averaged between 40 and 60 min after second injection for each hippocampal subregion. Dots indicate the mean; p-values and Cohen’s d are shown after FDR correction. Psilocybin significantly decreases the proportion of bursting units in CA1 (p = 0.044, d = −0.82), CA2 (p = 0.031, d = −0.90), and CA3 (p < 0.001, d = −1.44); DG and subicular complex show no significant changes. Ket + Psi produces no significant effects in any hippocampal subregion.

At the level of thalamic subregions, relative to saline, psilocybin produced opposing, nucleus-specific effects on burst rate (Figure 2c; for raw signal and absolute changes see Figure S3). The reticular nucleus showed a significant increase in burst rate (FDR-corrected p < 0.001, d = 0.82), with a comparably strong increase in the anterior thalamic nuclei (FDR-corrected p < 0.001, d = 0.76); both peaked at approximately 45–50 min post-injection, though the RT increase emerged earlier, reaching significance from ∼10 min onward, while the anterior increase was confined to the 40–50 min window. First-order somatomotor relay nuclei showed a smaller but significant increase (FDR-corrected p = 0.006, d = 0.52), concentrated in the same 45–50 min window. In contrast, first-order sensory geniculate nuclei showed a significant decrease in burst rate (FDR-corrected p = 0.049, d = −0.34), most consistent between 30–45 min, indicating that psilocybin suppressed relay burst output in geniculate nuclei while simultaneously driving burst-mode firing in RT and limbic/motor thalamus. Higher-order nuclei showed a nominal trend toward increased burst rate that did not survive FDR correction (uncorrected p = 0.05, d = 0.62), while intralaminar nuclei showed no significant change in burst rate. Critically, all three burst rate increases (RT, anterior, somatomotor) were abolished by ketanserin pretreatment (all FDR-corrected p > 0.85), indicating these effects were at least partially mediated by 5-HT2A receptors. The geniculate decrease could not be evaluated under ketanserin in either metric, as units in this region were sampled in only 2 of 8 Ket+Psi sessions, below our ≥3-session inclusion criterion. Burst fraction largely confirmed the burst rate pattern: under psilocybin neurons in the RT showed a significant increase (FDR-corrected p = 0.019, d = 0.57) while geniculate nuclei cells significantly decreased (FDR-corrected p = 0.019, d = −0.56), consistent in direction and magnitude with the burst rate findings above; as with burst rate, the RT increase was abolished by ketanserin pretreatment (FDR-corrected p = 0.853; Figure S4a). Anterior, higher-order, somatomotor, and intralaminar nuclei showed no significant burst fraction change. Burst size also increased significantly after psilocybin relative to saline in RT, anterior, and higher-order nuclei (FDR-corrected p = 0.024 for all three; d = 0.47, 0.32, and 0.20, respectively), while somatomotor, geniculate, and intralaminar nuclei showed no significant change (Figure S4a).

Within isocortex, burst fraction in prefrontal cortex decreased from baseline under saline injection in deep layers, while psilocybin maintained burst fraction near baseline, yielding a nominal drug–saline contrast in layer 5 (uncorrected p = 0.032, d = 0.24) and layer 6 (uncorrected p = 0.012, d = 0.35; Figure 2d). Neither effect survived FDR correction (FDR-corrected p = 0.177 and 0.130, respectively). Burst rate showed a similar, more extensive pattern: prefrontal cortex showed a nominal drug–saline contrast in layer 2/3 (uncorrected p = 0.028, d = 0.36), layer 5 (uncorrected p = 0.009, d = 0.22), and layer 6 (uncorrected p = 0.046, d = 0.21), consistent in direction with the burst fraction findings above (Figure S4b). Layer 4 of somatosensory cortex showed a separate nominal effect on burst rate, with psilocybin increasing burst rate relative to saline (uncorrected p = 0.024, d = 0.56). No other cortical area or layer showed a nominal drug–saline contrast in either metric. None of these burst rate effects survived FDR correction across the full set of cortical layers tested (all FDR-corrected p ≥ 0.099); we therefore report uncorrected p-values, shown in brackets in Figure 2d and Figure S4b, and interpret both the burst fraction and burst rate effects as trends rather than confirmed effects. Ketanserin pretreatment abolished the prefrontal contrast trend in both metrics (all uncorrected p > 0.31), paralleling the ketanserin effect observed in thalamus. Burst size was not significantly affected by either psilocybin or ketanserin administration in any cortical area or layer (all FDR-corrected p ≥ 0.645; Figure S4b).

In the hippocampus, psilocybin reduced the proportion of bursting units relative to baseline in CA1, CA2, and CA3 in the 40–60 min post-injection window (Figure 2e; for raw timecourse, see Figure S5), reaching significance relative to saline in all three subfields: CA3 (FDR-corrected p < 0.001, d = −1.44), CA2 (FDR-corrected p = 0.031, d = −0.90), and CA1 (FDR-corrected p = 0.044, d = −0.82). The dentate gyrus and subiculum showed no significant changes (FDR-corrected p = 0.112 and 0.317, respectively). Ketanserin pretreatment abolished the burst suppression in CA1 and CA3 (FDR-corrected p = 0.575 and 0.358, respectively), with no significant effects observed in any hippocampal subregion tested under Ket+Psi (Figure 2e, S5), consistent with at least partial 5-HT2A receptor mediation of psilocybin-induced hippocampal burst suppression. CA2 could not be evaluated under Ket+Psi, as units in this subregion were sampled in only 2 of 8 Ket+Psi sessions, below our ≥3-session inclusion criterion. Burst rate and burst size showed no significant changes in any hippocampal area under either psilocybin or Ket+Psi (all FDR-corrected p ≥ 0.149 and p ≥ 0.662, respectively; Figure S4c).

In the striatum (STR), psilocybin produced a significant increase in burst rate (p = 0.028, d = 0.27), an effect abolished by ketanserin pretreatment (p = 0.415), while burst fraction, mean burst size, and proportion of bursting units showed no significant changes (Figure S6).

Taken together, these findings revealed that psilocybin bidirectionally reconfigured burst firing in a region- and nucleus-specific manner: increasing burst rate in RT, anterior, and somatomotor thalamic nuclei and the striatum, suppressing it in geniculate relay nuclei and hippocampal CA1-CA3, and producing a trend toward preserved deep prefrontal layer bursting against a saline-driven suppression. Most of these effects were abolished by ketanserin pretreatment, indicating at least partial 5-HT2A receptor mediation. Mean burst size followed a distinct, more spatially restricted pattern: it increased selectively in reticular, anterior, and higher-order thalamic nuclei (Figure S4a) but was not significantly altered in isocortex or hippocampus (Figure S4b, c), indicating that psilocybin’s effect on the internal structure of bursts was largely confined to thalamus, while the cortical and hippocampal changes instead reflected altered burst frequency and prevalence rather than burst structure itself.

### Burst dynamics is dissociated from firing rate which is selectively decreased by psilocybin and further suppressed by ketanserin

To determine whether the burst changes described above were accompanied by changes in mean firing rate, we analyzed unit firing rate activity across all recorded brain regions and sessions for all three experimental conditions (Figure 3a). As with the burst analysis, firing rate is modulated by behavioral state (Stringer et al. 2019); we therefore likewise restricted this analysis to periods of rest (speed < 1 cm/s; see Methods) to allow for direct comparison between the two measures. Saline animals exhibited a gradual increase in firing rate over the course of the recording session, consistent with a time-dependent drift observed in head-fixed preparations in separate datasets in the absence of pharmacological manipulation (Figure S7); this drift likely reflects cumulative stress responses due to head-fixation (Juczewski et al. 2020) and/or mechanical effects of acute probe recordings. After the second injection, the psilocybin group diverged from saline, with modest drug-induced reductions superimposed on top of this slowly rising baseline (Figure 3a). The Ket+Psi group instead showed an immediate decrease in firing rate following the first (ketanserin) injection, consistent with the acute removal of tonic 5-HT2A-mediated excitatory drive, and after the second injection produced a more pronounced and sustained suppression than psilocybin alone, likewise superimposed on the rising baseline (Figure 3a, b).

**Figure 3.**
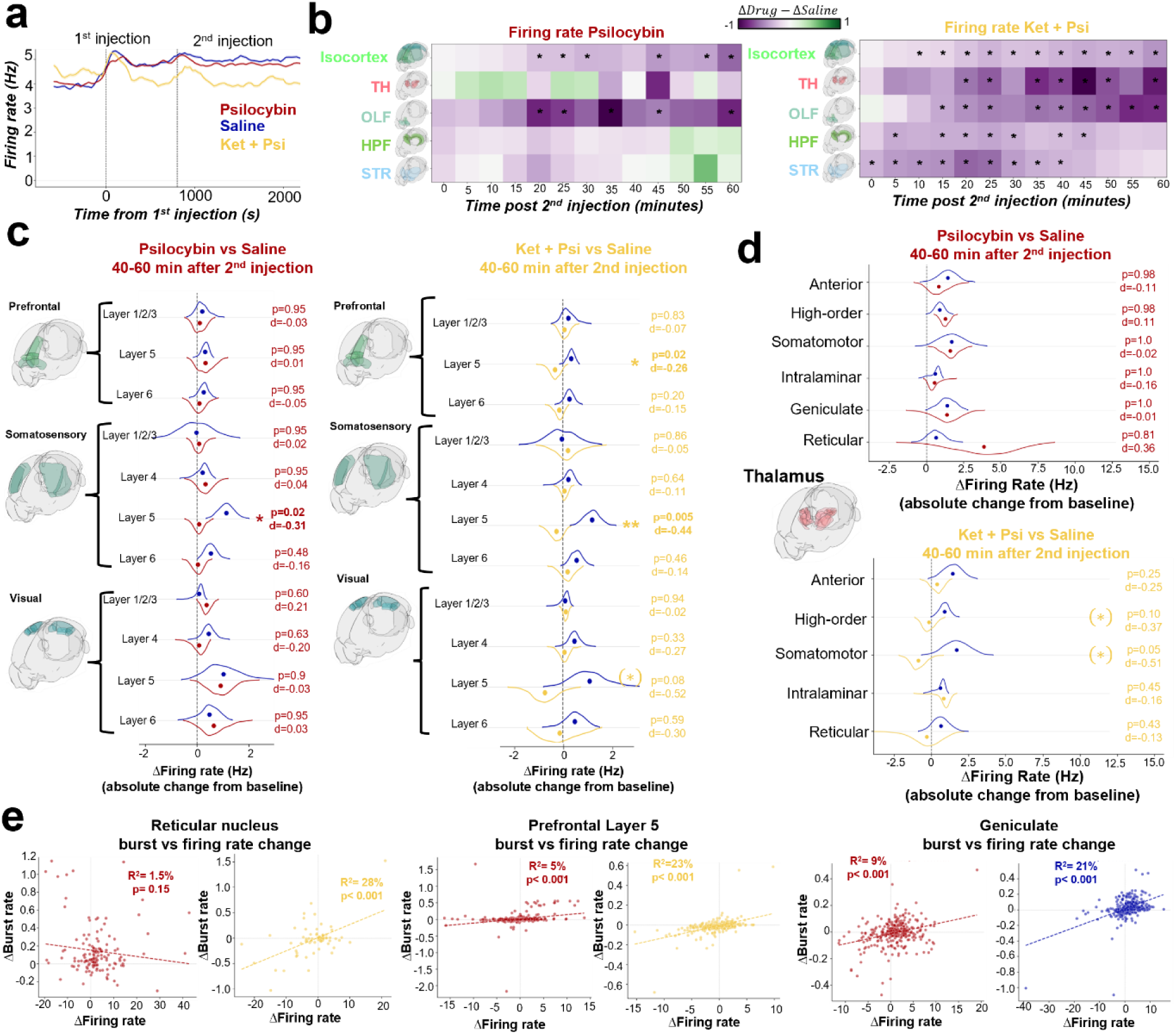
Firing rate is selectively decreased by psilocybin and significantly suppressed brain-wide by ketanserin, dissociated from psilocybin-induced burst changes. a, Mean firing rate of all recorded RS units from baseline to ∼40 min after first injection (at time 0 s) for psilocybin (red), saline (blue), and Ket + Psi (yellow) groups. Dashed vertical lines indicate the first and the average second i.p. injection timepoints following the design shown in Figure 1d. Note: the Ket + Psi group showed an overall decrease in firing rate immediately after the first injection, probably due to the acute blockade of tonic 5-HT2A receptor signaling, which normally sustains baseline cortical excitability. b, Heatmaps showing the difference in mean per-unit firing rate change (drug minus saline; absolute change from pre-injection baseline, in spikes/s) computed in 5-min bins after the second injection, across five major brain areas (Isocortex, Thalamus, Olfactory areas, Hippocampal formation, Striatum) for psilocybin (left) and Ket + Psi (right). Per-unit firing rate changes from baseline were computed at the single-unit level, averaged within each condition, and the saline group mean subtracted to isolate drug-specific effects. The diverging colormap indicates the direction of the drug–saline difference: green = drug above saline (facilitation), purple = drug below saline (suppression). Asterisks mark time bins where the drug condition differed significantly from saline (hierarchical bootstrap test, p < 0.05, uncorrected). Statistical comparisons were restricted to the first 60 min post-injection. Note that Ket + Psi induced a broad decrease in firing rate across all recorded brain areas, consistent with widespread 5-HT2A blockade reducing tonic cortical and subcortical excitability. c, Firing rate of RS units in isocortex subregions stratified by layer. Half-violin plots show the distribution of firing rate averaged between 40 and 60 min after the second injection for prefrontal (top), somatosensory (middle), and visual (bottom) cortex, stratified by layer. Dots indicate the mean; p-values and Cohen’s d are shown after FDR correction. Left column: psilocybin vs. saline. Right column: Ket + Psi vs. saline. Ket + Psi reduced firing rate in deep layers of prefrontal (layer 5: p = 0.02, d = −0.26) and somatosensory (layer 5: p = 0.005, d = −0.44) cortex, with a similar trend in visual cortex layer 5 that did not survive FDR correction (p = 0.08, d = −0.52). Psilocybin produced a significant decrease in firing rate in layer 5 of somatosensory cortex (p = 0.02, d = −0.31). d, As in c, for thalamic nuclei grouped by functional class (see Methods for details). Psilocybin did not induce significant changes in firing rate in any thalamic group. Ket + Psi produced a trend of decrease in higher-order (p = 0.10, d = −0.37) and first-order somatomotor (p = 0.05, d = −0.51) thalamic nuclei that did not survive FDR correction. e, Scatter plots of per-unit burst rate change vs. firing rate change in the reticular nucleus (left), prefrontal cortex layer 5 (middle, psilocybin and Ket + Psi), and geniculate nuclei (right, psilocybin and saline). R² and p-values are reported for each region and condition. In the reticular nucleus, the correlation was negligible and non-significant for psilocybin (R² = 1.5%, p = 0.15, red) but significant for Ket + Psi (R² = 28%, p < 0.001, yellow); the saline condition, shown in Figure S10, also showed a significant positive correlation (R² = 13%, p < 0.001). In prefrontal cortex layer 5, both psilocybin (R² = 5%, p < 0.001, red) and Ket + Psi (R² = 23%, p < 0.001, yellow) showed significant positive correlations; saline (Figure S10) showed a comparable significant positive correlation (R² = 13%, p < 0.001). The geniculate nuclei showed significant positive correlations for both psilocybin (R² = 9%, p < 0.001, red) and saline (R² = 21%, p < 0.001, blue), suggesting this relationship reflects an intrinsic property of geniculate neurons rather than a psilocybin-specific effect.

Regional heatmap analysis revealed that psilocybin-induced firing rate changes were spatially restricted, with sparse reductions mainly in olfactory areas and isocortex; no significant changes were detected in thalamus, hippocampal formation, or striatum (Figure 3b, left; Figure S8a). Ket+Psi produced a broader effect, reducing firing rates of RS units across all five major brain areas, with sustained suppression throughout the post-injection period in isocortex and olfactory areas, and significant but more transient effects in thalamus, hippocampal formation, and striatum (Figure 3b, right).

In the thalamus, psilocybin did not significantly alter firing rates in any thalamic group (Figure 3d, S8b). Ket+Psi produced a trend toward decreased firing rate in higher-order (uncorrected p = 0.042, d = −0.37) and first-order somatomotor (uncorrected p = 0.010, d = −0.51) thalamic nuclei; neither survived FDR correction (FDR-corrected p = 0.106 and 0.050, respectively) (Figure 3d, S8b).

At the cortical layer level, psilocybin produced a modest but significant decrease in firing rate in layer 5 of somatosensory cortex (FDR-corrected p = 0.022, d = −0.31; Figure 3c), the only cortical layer to survive FDR correction under psilocybin. No other cortical area or layer, including deep layers of prefrontal and visual cortex, showed a significant change in firing rate under psilocybin. Ket+Psi produced significant decreases in layer 5 of prefrontal cortex (FDR-corrected p = 0.022, d = −0.26) and layer 5 of somatosensory cortex (FDR-corrected p = 0.005, d = −0.44), with a non-significant trend toward suppression in layer 5 of visual cortex (uncorrected p = 0.025, d = −0.52; did not survive FDR correction, FDR-corrected p = 0.081), consistent with its broader suppressive profile in isocortex (Figure 3b, c, S8c). Layer 6 of prefrontal and somatosensory cortex did not reach significance under either drug condition.

The firing rate reductions under psilocybin were specific to RS units. Fast-spiking (FS) units showed no significant changes in firing rate under psilocybin across any brain region, whereas Ket+Psi produced a significant decrease in FS unit firing rate across isocortex, thalamus, hippocampal formation, and striatum, with olfactory areas notably unaffected (Figure S9).

Critically, in the reticular nucleus, the burst rate increase was dissociated from firing rate: per-unit burst rate changes were uncorrelated with firing rate changes specifically under psilocybin (R² = 0.02, non-significant) - the condition in which the burst rate increase occurred - even though a significant positive correlation between the two measures was present in both saline (R² = 0.14) and Ket+Psi (R² = 0.28) sessions (Figure 3e), indicating that psilocybin’s effect on RT burst rate could not be explained by a simple change in firing rate. In contrast, prefrontal cortex layer 5 and the geniculate nuclei showed significant positive correlations between per-unit firing rate and burst rate changes across all conditions tested (prefrontal layer 5: R² = 0.05–0.24 across psilocybin, saline, and Ket+Psi; geniculate: R² = 0.10–0.21 across psilocybin and saline; Ket+Psi geniculate data were excluded from analysis, as units in this region were sampled in only 2 of 8 Ket+Psi sessions, below our ≥3-session inclusion criterion), indicating that in these regions, unlike RT, burst rate and firing rate remained coupled properties of individual units regardless of drug condition (Figure S10). Together, these results demonstrated that psilocybin’s reconfiguration of burst dynamics was selectively decoupled from firing rate in the reticular nucleus specifically, while in prefrontal cortex and geniculate thalamus the two measures remained intrinsically linked.

### Psilocybin suppresses cortical high-gamma and resting-state alpha power, an effect not abolished by ketanserin pretreatment

To assess whether psilocybin-induced changes in single-unit activity were reflected at the mesoscale level, we extracted spectral power changes relative to baseline across six frequency bands (delta [0.1–4 Hz], theta [4–8 Hz], alpha [8–13 Hz], beta [13–35 Hz], gamma [35–55 Hz], and high gamma [65–100 Hz]) from the EEG and LFP signals (Figure 4a; see Methods).

**Figure 4.**
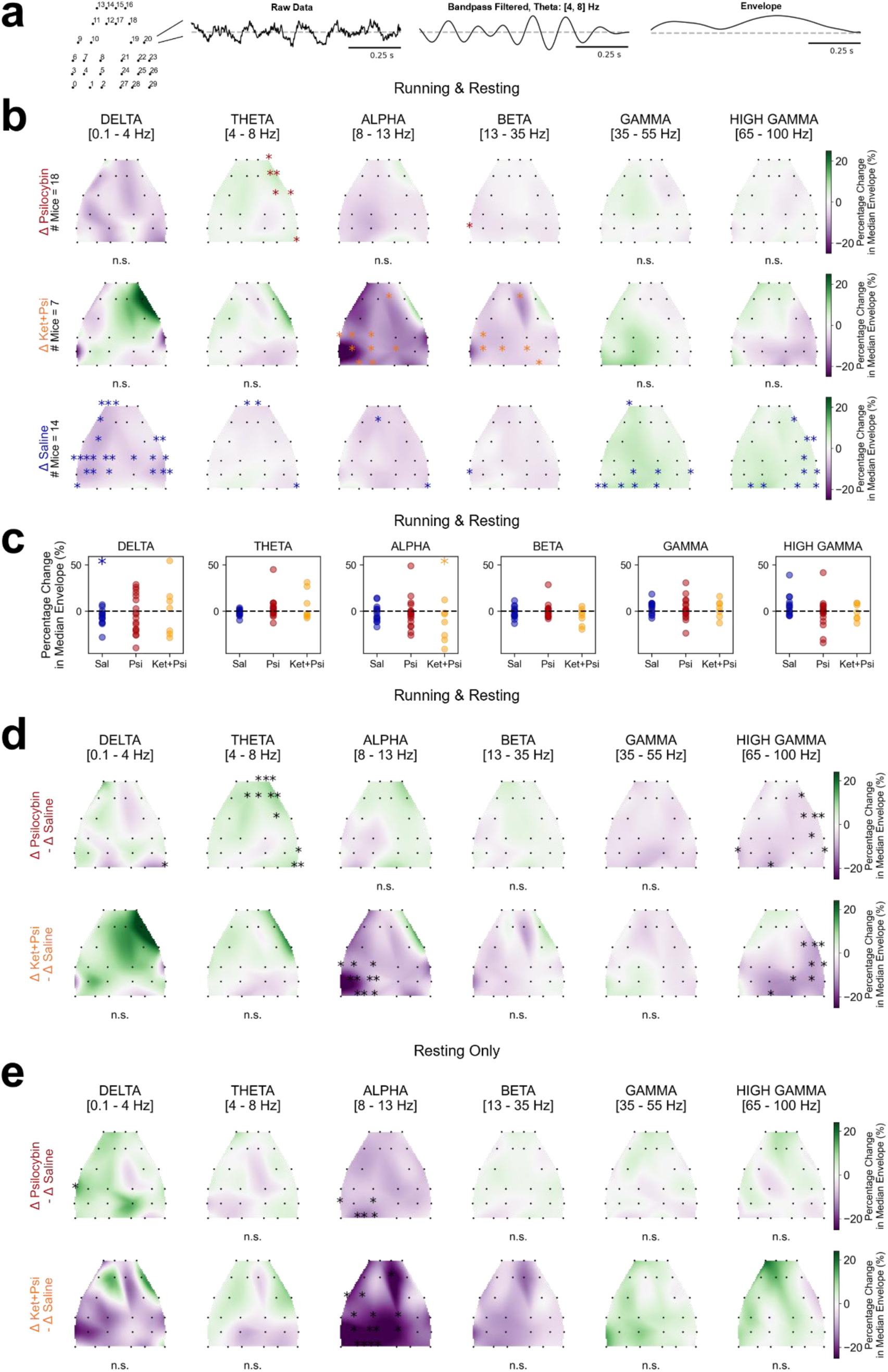
Psilocybin induces a decrease in both high-gamma and alpha EEG power, particularly evident during resting. Neither effect was abolished by ketanserin pretreatment. a, Illustration of the power analysis used to quantify changes in oscillatory EEG dynamics across frequency bands (see Methods for details). The figure shows the steps involved in calculating theta-band power from a one-second EEG segment of one of the electrodes shown in the schematics. b, Topographic plots of the median percentage change in power from baseline across all behavioral states (locomotion and rest combined) for psilocybin (top row, n=18 mice), Ket + Psi (middle row, n=7), and saline (bottom row, n=14) conditions, for each frequency band (columns) and EEG channel (black dots). Statistical significance at each channel was assessed using a Wilcoxon ranked-sum test; significant channels are marked with asterisks. c, Scatter plots of median percentage change in power from baseline averaged across channels, for saline (blue, n=14), psilocybin (red, n=18 mice), and Ket + Psi (yellow, n=7) conditions across each frequency band. Colored asterisks indicate a significant difference from baseline within each condition (Wilcoxon ranked-sum test). d, Topographic plots of the median percentage difference in power change between drug and saline conditions, for psilocybin minus saline (top row) and Ket + Psi minus saline (bottom row), across all behavioral states. Statistical significance at each channel was assessed using a Mann-Whitney U test; significant channels are marked with asterisks. e, Same as d, but restricted to resting periods only. Statistical results for each channel are reported in supplementary file Stats_EEG.xlsx.

Topographic plots of the median percentage change in EEG power from baseline across all behavioral states revealed an increase in gamma and high-gamma power and a decrease in delta power in saline animals distributed across multiple channels, not visible in psilocybin animals (Figure 4b, c; statistical details in supplementary file Stats_EEG.xlsx). Direct comparison of power changes between psilocybin and saline conditions confirmed a significant decrease in the high-gamma band in psilocybin animals (Figure 4d; supplementary file Stats_EEG.xlsx), consistent with the overall decrease in firing rate observed at the single-unit level. Compared to saline, an increase of theta power was visible after psilocybin (Figure 4d; supplementary file Stats_EEG.xlsx).

When the analysis was restricted to resting periods only, epochs during which the mouse was stationary across the full 10-second window (speed <1 cm/s), a significant decrease in alpha power emerged in the psilocybin condition for some of the occipital EEG channels. This reduction was not apparent when all behavioral states were combined (Figure 4e, S11; statistical details in supplementary file Stats_EEG.xlsx). This suggested that the alpha power decrease was partially obscured by locomotion-related activity and became detectable specifically during periods of quiet rest. Critically, Ket+Psi animals showed a pattern of EEG and LFP power changes closely mirroring that of the psilocybin group and even amplified, with significant decreases in high-gamma power across all behavioral states and a significant alpha power decrease during resting periods (Figure 4d, e, S11, S12, statistical details in supplementary file Stats_LFP.xlsx), indicating that these spectral effects were not abolished by ketanserin pretreatment, and were, if anything, amplified by it.

## DISCUSSION

We provide a brain-wide single-unit characterization of psilocybin’s acute effects across the CSTC circuit, demonstrating that a single administration produced dissociable effects on neural activity through distinct serotonergic mechanisms: a selective, region-specific modulation of firing rates that was not reversed by 5-HT2A blockade, and a selective 5-HT2A-dependent reconfiguration of burst coding whose regional specificity was most pronounced at the thalamic gate, pointing to altered thalamic filtering as a core circuit mechanism of psilocybin’s acute action.

Psilocybin bidirectionally reconfigured thalamic burst coding in a pattern consistent with altered thalamic gating. Among the strongest effects in the entire dataset were marked increases in burst rate in the reticular nucleus (RT) and the anterior thalamus, with a smaller but significant increase also observed in first-order somatomotor relay thalamic nuclei. This effect was abolished by ketanserin pretreatment. Instead, the geniculate nuclei showed the opposite pattern: a significant psilocybin-induced decrease in both burst rate and burst fraction. These effects are anatomically consistent with 5-HT2A receptor distribution: RT neurons express high levels of both 5-HT2A and 5-HT1A receptors throughout the nucleus, and RT receives dense serotonergic innervation primarily from the dorsal raphe (Rodriguez et al. 2011). The anterior thalamic nuclei likewise receive unusually dense direct raphe serotonergic innervation, denser than most other thalamic territories (Gonzalo-Ruiz, Lieberman, and Sanz-Anquela 1995; Vertes, Linley, and Hoover 2010). Consistent with our findings, tonic serotonergic input increases burst-firing mode in RT neurons of anesthetized rats via 5-HT1A receptor activation (Barrientos et al. 2022). Nonetheless, the abolition of the RT burst increase by ketanserin in our data is consistent with at least partial 5-HT2A-receptor mediation of this shift in RT burst dynamics in awake, behaving mice. Serotonin’s opposite effects on RT/anterior/somatomotor versus geniculate burst firing are consistent with a distinction between postsynaptic and presynaptic loci of action: RT, anterior, and somatomotor thalamus share the postsynaptic mechanism described above, whereas serotonin acts presynaptically at the retinogeniculate synapse to suppress glutamate release from retinal ganglion cell terminals (Seeburg, Liu, and Chen 2004), reducing the driving afferent input that normally triggers relay-cell bursts rather than altering the postsynaptic membrane state directly. The RT burst-rate increase was uncorrelated with firing-rate change under psilocybin, ruling out a simple gain-change account; because the two measures were correlated under saline and Ket+Psi, this dissociation is specific to the psilocybin condition rather than a fixed property of RT. In the geniculate, firing rate and burst rate were coupled under both saline and psilocybin, but because psilocybin did not alter mean geniculate firing rate, the burst-rate decrease cannot be a downstream consequence of a rate change and instead reflects a direct action on burst dynamics, consistent with the presynaptic mechanism described above. Ketanserin pretreatment produced a trend toward decreased firing rate in higher-order and somatomotor nuclei, paralleling the suppression in deep prefrontal and somatosensory layers and consistent with reduced corticothalamic drive to these reciprocally connected relay nuclei. Together, these findings provided single-unit mechanistic grounding for the thalamic filtering changes observed in human neuroimaging under serotonergic psychedelics (Preller et al. 2019; Preller et al. 2018; Muller et al. 2017) and the proposed role of altered CSTC dynamics in serotonergic psychedelic phenomenology (Geyer and Vollenweider 2008; Llinas et al. 1998). More broadly, thalamic nuclei are increasingly recognized as active contributors to predictive processing rather than passive relays: the higher-order visual nucleus LP, for example, conveys the discrepancy between self-generated and external visual motion to cortex (Roth et al. 2016). Crick’s searchlight hypothesis proposed that RT’s inhibitory output does not merely suppress thalamic relay cells, but switches them into burst mode. That is, bursting signals which thalamocortical channel is transiently amplified, gating what reaches cortex (Crick 1984). Our finding that psilocybin drove RT toward burst mode while simultaneously suppressing burst output in the first-order geniculate relay is consistent with a pharmacological co-optation of this gating mechanism: rather than being recruited by corticothalamic feedback, RT’s burst-mode shift was instead driven by 5-HT2A activation, decoupling thalamic gating from top-down cortical control. This offers a circuit-level account of the rebalancing between bottom-up sensory throughput and top-down predictive control that is central to several current models of psychedelic action (Carhart-Harris and Friston 2019; Corlett et al. 2019), by identifying the thalamic reticular nucleus as a site where this balance can be shifted independently of cortex.

Although not a canonical CSTC node, the hippocampus has been proposed as a target of psilocybin’s effects on trauma and mood (Carhart-Harris et al. 2017; Catlow et al. 2013). Psilocybin produced a significant, ketanserin-sensitive reduction in CA3 burst participation, consistent with 5-HT2A dependence. CA3 is well suited to this effect: its dense recurrent collaterals form an auto-associative attractor network specialized for pattern completion (Nakazawa et al. 2002; van Strien, Cappaert, and Witter 2009; Rolls 2007), its pyramidal neurons are among the most intrinsically burst-prone in the brain (Roy and Narayanan 2023; Raus Balind et al. 2019), and 5-HT2A receptors are densely expressed on their somata and dendrites (Pompeiano, Palacios, and Mengod 1994). By dampening CA3 bursting, psilocybin may reduce the recurrent drive sustaining attractor states that, when pathologically reinforced, underlie fear overgeneralization and post-traumatic stress disorders (Bains, Longacher, and Staley 1999; Kheirbek et al. 2012) - a candidate acute mechanism for disrupting maladaptive memory traces. As in the thalamus, firing rate was unaffected.

Head-fixation produces a pronounced acute stress response, with corticosterone reaching up to 9-fold above baseline levels (Juczewski et al. 2020); stress-induced catecholamine release preferentially targets prefrontal layer 5 pyramidal neurons, where α1-adrenergic and D1 signaling opens potassium channels that weaken synaptic efficacy and suppress burst firing (Arnsten 2015, 2009; Guo et al. 2014). Consistent with this, burst fraction in deep prefrontal layers decreased from baseline under saline, while psilocybin maintained burst fraction near baseline, yielding a nominal drug–saline contrast in layer 5 and layer 6 that did not survive FDR correction. Somatosensory and visual cortices showed no comparable effect. This trend was abolished by ketanserin pretreatment, consistent with at least partial 5-HT2A dependence, in line with genetic evidence that 5-HT2A receptors were required for psilocybin’s structural and behavioral effects in frontal cortex (Shao et al. 2025). Although modest at the population level, this pattern echoed prior work on psilocybin-evoked structural plasticity in prefrontal layer 5: rapid, persistent dendritic spine growth (Shao et al. 2021) that is activity-dependent (Jiang et al. 2026) and requires specifically the subcortically-projecting PT neurons of deep layer 5, sparing neighboring IT neurons (Shao et al. 2025). PT neurons are intrinsically more burst-prone than IT neurons owing to distinct dendritic calcium electrogenesis (Larkum 2013; Wang and McCormick 1993; Kasper et al. 1994), and their burst firing is itself suppressed by α1-adrenergic activation (Wang and McCormick 1993), converging with the stress pathway described above (Arnsten 2015, 2009). Because our RS classification pools PT and IT neurons, this mismatch, together with the non-specific firing-rate drift we attribute to prolonged head-fixation and/or probe settling, may dilute a genuinely PT-specific effect, offering one explanation for the weak population-level signal. Burst firing is more closely linked to plasticity induction than isolated spikes, through supralinear dendritic calcium events and BDNF/TrkB signaling (Lisman 1997; Magee and Johnston 1997; Harris et al. 2001; Pike et al. 1999; Ly et al. 2018; Larkum 2013); because psilocybin’s synaptic rewiring is selective for active synapses (Jiang et al. 2026), firing pattern rather than rate may help determine which connections are strengthened. Consistent with this, psilocybin significantly decreased firing rate in deep somatosensory cortex, confirming other studies (Purple et al. 2025; Skyberg et al. 2025; De Filippo et al. Under submission). This effect was not reversed by ketanserin pretreatment, consistent with a 5-HT2A-independent mechanism of intrinsic excitability suppression (Ekins et al. 2023).

Critically, these electrophysiological effects were not secondary to locomotor changes: all analyses were restricted to resting epochs, and psilocybin did not alter locomotion speed, consistent with prior reports (Fadahunsi et al. 2022; Lu et al. 2025; Purple et al. 2025). Subtle autonomic responses such as pupil dilation described in humans (Carbonaro et al. 2018; Passie et al. 2002) were detectable through within-subject comparisons with mixed-effects modelling (Figure S1), underscoring the importance of paired experimental designs in psychedelic electrophysiology as demonstrated in our other independent study (De Filippo et al. Under submission). Both the pupil and locomotor responses and the concurrent EEG decrease in alpha and high-gamma power were not abolished by ketanserin pretreatment. Strikingly, ketanserin pretreatment induced a broad, previously unreported suppression of firing rates across brain regions, affecting both regular- and fast-spiking units, a pattern consistent with removal of the tonic excitatory drive sustained by endogenous serotonin at 5-HT2A/2C receptors under baseline conditions.

Our results revealed a fundamental dissociation at the heart of psilocybin’s neural actions: a modest firing-rate suppression largely confined to somatosensory cortex, and a broader EEG power suppression, neither of which was abolished by ketanserin pretreatment, whereas 5-HT2A-dependent modulation of burst coding represented a targeted reorganization of neural computation across the CSTC circuit. These burst changes were spatially precise yet circuit-level coherent, spanning reticular, anterior, somatomotor, and geniculate thalamus, striatum, hippocampal CA1–CA3, and deep prefrontal layers, nodes that together gate sensory information, relay cortical context, and encode associative memories. The psychedelic state, these data suggest, is not simply a matter of how much neurons fire, but of how they fire, and it is this reconfiguration of coding mode, rather than net excitation, that may underlie the acute phenomenology of psilocybin and possibly its lasting therapeutic effects.

## METHODS

Experimental procedures followed closely those described in our previous studies (Claar et al. 2023; Russo et al. 2025). The methods are summarized below, and additional details are provided for novel procedures.

### Mice

Male and female C57BL/6J wild-type mice (n=40) were purchased from Jackson Laboratories (JAX stock #000664) on postnatal day 28–35. Mice were housed in the Allen Institute animal facility and used in accordance with protocols approved by the Allen Institute’s Institutional Animal Care and Use Committee (protocol 2212). All mice were housed under controlled room temperatures (20–22°C) and humidity (30–70%), maintained on a reverse 12-hour light cycle, and single-housed following surgery with ad libitum access to food and water.

### Surgical procedures and Habituation

#### Headframe and EEG Implantation Surgery

Prior to experimental procedures, all mice were implanted with a titanium headframe to facilitate head fixation in vivo (surgical procedures adapted from (Groblewski et al. 2020). An EEG array was implanted on the skull during the same surgery. Surgery took place when mice were 4–11 weeks old. Detailed surgical procedures are available on Protocols.io (Marks, Rembado, and Claar). Briefly, pre-operative injections of dexamethasone (3–4 mg/kg, IM) and ceftriaxone (100–125 mg/kg, SC) were administered 1–3 hours prior to surgery. Mice were deeply anesthetized with isoflurane (5%) in an induction chamber, then placed on a stereotaxic rig (Model 1900, Kopf Instruments, Tujunga, CA) and maintained at a surgical level of anesthesia with isoflurane (1.5–2.5%) via nose cone. Breathing was monitored throughout, and body temperature was maintained at 37.5°C using a feedback-controlled heating pad (TC-1000, CWE Inc.). Ocular lubricant (Systane, Alcon Inc., Geneva, Switzerland) was applied to both eyes to prevent desiccation. The skull was exposed and leveled with reference to bregma. Three skull screws were implanted above the cerebellum to serve as electrical ground and reference for electrophysiological recordings. A 30-channel EEG array was secured to the skull surface using silicone bio-adhesive (Kwik-Cast, World Precision Instruments). A custom titanium headframe was secured to the skull using white C&B Metabond (Parkell Inc., Edgewood, NY), and the remaining exposed skull was covered with the same cement. Post-operatively, mice received a subcutaneous injection of lactated Ringer’s solution (up to 1 mL) and were placed on a heating pad to recover. All animals received analgesics and antibiotics for two days post-surgery.

#### Habituation

Five days after headframe implantation surgery, all mice began a 3-week training schedule. Researchers first habituated mice to handling for 2 days; on all subsequent days, mice were head-fixed on a running wheel using two screws attached to the headframe clamp (Groblewski et al. 2020). Session durations increased progressively from 5 to 90 min. During the final week of training, mice received two sham intraperitoneal (IP) injections per session to habituate them to the injection procedure in the head-fixed state; no liquid was administered during sham injections.

All training and experimental sessions were conducted during the dark cycle. A subset of mice (n=11) underwent a locked-wheel training protocol, in which the running wheel was locked during the habituation period to reduce locomotion during subsequent experimental sessions.

#### Burr Hole Craniotomy Surgery

One day prior to the first recording session, mice underwent surgery to create four burr hole craniotomies (0.5–1 mm diameter) in the left hemisphere for subsequent Neuropixels probe insertion. Detailed procedures are available on Protocols.io (Marks, Rembado, and Claar). Briefly, mice were anesthetized and secured in the stereotaxic frame using the headframe clamp. Small openings (1–2 mm diameter) were made in the dental cement at the planned probe insertion sites, the Kwik-Cast underneath was removed with a scalpel, and a small hole was cut in the polyimide layer of the EEG array. Burr holes (0.5–1.5 mm diameter) were then drilled in the skull. Exposed brain tissue was covered with artificial cerebrospinal fluid (ACSF) and Kwik-Cast. A plastic reservoir was cemented around the perimeter of the craniotomies to hold ACSF during recordings. Post-operatively, mice received a subcutaneous injection of lactated Ringer’s solution (up to 1 mL) and were placed on a heating pad to recover. All animals received analgesics and antibiotics for two days post-surgery.

### Experimental timeline

After completing three weeks of habituation, mice underwent one recording session per day. Each session followed one of three experimental conditions depending on the assigned group: saline control, psilocybin, or ketanserin+psilocybin (see IP injections section below for drug administration details).

After signal quality control (see below) a total of 35 mice were used across all experiments. Across 47 recording sessions, 17 sessions were saline control sessions in which mice received two saline injections and are referred to as the saline group. 22 sessions were psilocybin sessions in which mice received a saline injection followed by psilocybin and are referred to as the psilocybin group. 8 sessions were ketanserin+psilocybin sessions in which mice received a ketanserin injection followed by psilocybin and are referred to as the Ket+Psi group.

To enable within-subject comparisons, a subset of mice underwent both a saline session and a psilocybin session on consecutive days (paired saline–psilocybin group, n=7 mice). Similarly, a separate subset of mice underwent both a saline session and a Ket+Psi session on consecutive days (paired saline–Ket+Psi group, n=5 mice). EEG and LFP analyses drew from a separate, partially overlapping set of sessions, since each signal modality was subject to its own independent quality-control criteria (see EEG quality control and pre-processing, and Neuropixels signals pre-processing); session counts for those analyses are reported in their respective Methods subsections and figure legends.

During each recording session, mice were placed on the running wheel and head-fixed using the headframe clamp. A thin layer of Kwik-Cast was removed from the craniotomies and the plastic reservoir was filled with ACSF to maintain brain tissue hydration. A plastic cone was positioned above the mouse to prevent the tail from contacting the brain surface or probes. A grey screen was placed in front of the mouse and a black curtain was lowered to keep the animal in darkness throughout the session.

A protocol was developed to administer intraperitoneal (IP) injections while animals were head-fixed on the wheel. The researcher scruffed the loose skin over the mouse’s back to immobilize the hindlimbs, and the injection was administered into the lower right quadrant of the peritoneal cavity. In psilocybin sessions, mice received a first injection of sterile saline (0.01 mL/g, IP) followed 10–12 min later by psilocybin (1 mg/kg, IP; Usona Institute). In Ket+Psi sessions, mice received a first injection of ketanserin (1 mg/kg, IP) to block 5-HT2A receptors, followed 10–12 min later by psilocybin (1 mg/kg, IP; Usona Institute). In saline control sessions, two injections of sterile saline (0.01 mL/g, IP) were administered 10–12 min apart. To administer ketanserin (Sigma-Aldrich, #S006) we first dissolved it in a solution of 10% DMSO in saline to a concentration of 0.1 mg/ml. On the day of experiment, the mouse was given a dose of 1 mg/kg. The dilution was stored at −20 °C for up to 1 month.

### EEG and Neuropixels recording and cortical electrical stimulation

The implanted 30-electrode EEG array was connected to a 32-channel headstage (RHD 32ch, Intan Technologies, Los Angeles, CA) controlled by an Open Ephys acquisition board (Siegle et al. 2021; Claar et al. 2023; Russo et al. 2025). EEG signals were sampled at 2.5 kHz with a low-frequency cutoff of 0.1 Hz and digitized at 16-bit resolution. The EEG array consisted of 30 circular platinum electrodes (diameter: 500 μm; impedance: 0.01 MΩ as measured by the manufacturer). One skull screw implanted over the cerebellum served as electrical ground and a separate screw served as the reference electrode.

Up to four Neuropixels 1.0 probes were inserted per session, targeting motor (MO), somatosensory (SS), visual (VIS), and thalamic regions of the left hemisphere. Each probe was inserted individually using a three-axis micromanipulator (New Scale Technologies, Victor, NY) at a rate of 200 μm/min to a depth of up to 3.5 mm. Probes were allowed to settle for 10–15 min before recording commenced. Signals from each recording site were split in hardware into a spike band (30 kHz sampling rate, 300 Hz high-pass filter, 500× gain) and an LFP band (2.5 kHz sampling rate, 300 Hz low-pass filter, 250× gain), and acquired using the Open Ephys GUI (Siegle et al. 2017). Each probe was connected to a dedicated PXIe card inside a National Instruments chassis. The reference connection on each Neuropixels probe was permanently soldered to an insulated silver wire and connected to one of the skull screws implanted over the cerebellum. All recordings used external reference and ground configuration using the cerebellum skull screws. Some animals received electrical stimulation delivered to the cortex via a custom bipolar platinum-iridium stereotrode (Microprobes for Life Science, Gaithersburg, MD) consisting of two parallel monopolar electrodes (50 kΩ impedance, 300 μm vertical tip offset) inserted to target layers 5/6 of the left secondary motor cortex (MOs; 1.4 ± 0.24 mm below the brain surface). Each stimulation block consisted of up to 80 biphasic, charge-balanced, cathodic-first current pulses (200 μs per phase, 6.5–7.5 s jittered inter-stimulus interval) delivered at three current intensities selected individually for each animal: the maximum intensity was the largest current below 100 μA that evoked a visible EEG response without eliciting visible muscle twitches; the minimum intensity was the smallest current evoking a visible EEG response; and the intermediate intensity was the average of the maximum and minimum (High: 66.15±12.11 μA; Intermediate: 47.35±13.30 μA; Low: 30.59±15.80 μA). Prior to insertion, all Neuropixels probes and the stimulating electrode were coated with a fluorescent dye (Vybrant DiI/DiO, ThermoFisher Scientific, Waltham, MA) by repeated immersion and slow withdrawal to allow drying.

### Recording timeline

Data were collected for 86–170 min per session. The experimental timeline followed one of two protocols: (1) 10-15 minute baseline period, one block of electrical stimulation, first saline or ketanserin injection, 10–12 min of spontaneous activity, second injection (saline or psilocybin), 15 min of spontaneous activity, one block of electrical stimulation; or (2) 30-45 minute spontaneous activity, 1^st^ saline or ketanserin injection, 10-12 minute spontaneous activity, second injection (saline or psilocybin), 30-90 min spontaneous activity.

### Behavioral Data

Videos of the eye and body were acquired at 30 Hz and 60 Hz, respectively. The angular position of the running wheel was acquired by a dedicated computer with a National Instruments card sampling digital inputs at 100 kHz, which served as the master clock. A 32-bit digital barcode was sent via an Arduino Uno (DEV-11021, SparkFun Electronics, Niwot, CO) every 30 s to synchronize all recording devices (Siegle et al. 2021).

### Ex vivo imaging and localization of electrodes

To localize the probes, we followed the procedure we extensively described in Claar et al., 2023 (Claar et al. 2023). Briefly, after the final recording session, mice were deeply anesthetized (5% isoflurane) and transcardially perfused with 4% paraformaldehyde. Brains were post-fixed in 4% paraformaldehyde for 48 hours, rinsed with 1× PBS, and stored at 4°C in PBS with 0.1% sodium azide. Whole brains were imaged using serial two-photon tomography (Ragan et al. 2012; Oh et al. 2014). Images were aligned to the Allen Mouse Brain Common Coordinate Framework version 3 (CCFv3) following the process described by Oh et al. (2014) (Oh et al. 2014). Fluorescent tracks corresponding to Neuropixels probe and stimulation electrode locations were manually identified in 10 μm affine-aligned space and warped to 25 μm CCF space using custom software (Tissuecyte_Annotation). The same software was used to align major structural boundaries along each probe track with the corresponding physiological data (Liu et al. 2021; Siegle et al. 2021), and each recording channel and associated units were assigned to a unique CCFv3 brain structure.

### EEG quality control and pre-processing

Before each experiment, EEG signal quality was assessed by exposing the animal to visual flashes and evaluating the signal-to-noise ratio of evoked responses. Animals with low signal-to-noise ratio, excessive 60 Hz noise, or large movement artifacts were excluded (n=8 sessions). Experimental EEG data were preprocessed as follows: recordings were visually inspected to identify artifact-contaminated electrodes, which were excluded from further analysis (mean ± SD electrodes excluded per subject: 3.9 ± 3.7 out of 30). EEG signals from all remaining electrodes were downsampled to 500 Hz.

### Neuropixels signals pre-processing

Neuropixels action potential (AP) raw signals were artifact masked by replacing 0–2ms stimulation artifact window with the raw signal from –2–0ms before being pre-processed and spike-sorted using Kilosort 2.0 (Stringer et al. 2019) as described by Siegle, Jia et al. (Siegle et al. 2021). After spike sorting, any spikes that occurred during the artifact window (0 to +2 ms from stimulus onset) were removed from further analysis. High quality units were identified for further analysis using metrics described by Siegle, Jia et al (Siegle et al. 2021). We classified cortical regular spiking (**RS**) and fast spiking (**FS**) neurons (putative pyramidal and inhibitory neurons, respectively) based on their spike waveform duration (RS duration > 400 μs; FS duration ≤ 400 μs) (Bartho et al. 2004; Bruno and Simons 2002; Bortone, Olsen, and Scanziani 2014; Niell and Stryker 2008; Sirota et al. 2008). Similarly, thalamic units were classified as putative relay neurons if their spike width was above 400 μs (Guo et al. 2017; Bartho et al. 2014; Huo, Chen, and Guo 2020). This classification was not applied to the striatum and the reticular nucleus of the thalamus where all the units were considered for further analyses.

Local field potential (LFP) signals were visually inspected to identify artifact-contaminated recordings, which were excluded from further analysis (6 excluded out of 47 total recordings). Remaining LFP recordings were downsampled to 500 Hz. For each probe, channels located in cortical areas were selected for further analysis.

### Neuroanatomical classification and region groupings

Units were assigned to brain regions using the Allen Mouse Brain Common Coordinate Framework version 3 (CCFv3). Region identity was resolved by walking the CCFv3 structure hierarchy to assign each unit to a major region (isocortex, hippocampal formation, olfactory areas, striatum, or thalamus) and, where applicable, to anatomical subregions and functional groups. For isocortical analyses, individual areas were grouped into four cortical categories. The prefrontal group comprised motor cortex (MO), orbital cortex (ORB), anterior cingulate cortex (ACA), prelimbic cortex (PL), infralimbic cortex (ILA), and frontal pole (FRP). The somatosensory group comprised all subdivisions of the primary and secondary somatosensory cortex (SS). The visual group comprised all subdivisions of the primary and higher visual cortical areas (VIS). The retrosplenial cortex (RSP) was analyzed separately and was not included in any of the above groups.

Thalamic nuclei were assigned to six functional groups based on established anatomical and physiological criteria (Sherman and Guillery 2006). The Anterior group included the anterodorsal (AD), anteroventral (AV), anteromedial (AMd, AMv), laterodorsal (LD), and interanterodorsal (IAD) nuclei (Nelson 2021; Hintiryan et al. 2025). The Higher-order group included mediodorsal (MD), lateral posterior (LP), posterior (PO), posterior triangular (PoT), and suprageniculate (SGN) nuclei (Sherman 2007; Halassa and Kastner 2017; Leow et al. 2022). The First-order Somatomotor group included the ventral anterolateral (VAL), ventral posterolateral (VPL), and ventral posteromedial (VPM) nuclei (Harris et al. 2019). The Intralaminar group included the central lateral (CL), central medial (CM), paracentral (PCN), parafascicular (PF), and subparafascicular parvicellular (SPFp) nuclei (Van der Werf, Witter, and Groenewegen 2002; Kumar, Scheffler, and Grodd 2023). The First-order Sensory Geniculate group included all subdivisions of the lateral geniculate nucleus (LGd, LGv, LGco, LGip, LGsh, IGL) and medial geniculate nucleus (MGd, MGm, MGv) (Sherman 2007; Harris et al. 2019). The Reticular nucleus (RT) was treated as a separate group in all analyses, given its unique GABAergic identity and functional role as a thalamic gating structure.

Within the hippocampal formation, units were assigned to CA1, CA2, CA3, and dentate gyrus (DG) based on their CCFv3 location. The subiculum (SUB) and prosubiculum (ProS) were merged into a single subicular complex group, as their anatomical boundary is difficult to resolve with Neuropixels probes and their functional properties are closely related.

### Data Analysis

#### Locomotion and Pupillometry

Locomotion speed was derived from the angular velocity of the running wheel, sampled at 100 kHz and converted to linear speed (cm/s). For population-level comparisons, mean locomotion speed was computed across the post-injection analysis window (1800–2400 s after the second injection) and compared to the mean speed during the baseline period pre-first injection for each session. Speed distributions were also computed for the same baseline and post-injection windows across saline, psilocybin, and Ket+Psi conditions.

Pupil size was extracted from videos of the right eye recorded during each experimental session using an Individual Model pipeline developed in-house based on DeepLabCut (Mathis et al. 2018) and described in detail in Seyfourian et al. 2026 (Seyfourian et al. 2026). Briefly, the pipeline fits an ellipse to the pupil boundary for each video frame, and the largest radius of the fitted ellipse was extracted over time to provide an estimate of pupil radius in pixels. The pupil radius (pixels) was normalized to the session baseline (pre-first injection) median during resting for all further analyses.

#### Burst Analysis

To characterize psilocybin-induced changes in neuronal firing patterns, we analyzed the bursting dynamics of units across brain regions and experimental conditions. Two distinct burst detection algorithms were applied depending on brain region (Figure 1f). For thalamic units, bursts were detected using canonical low-threshold calcium spike (LTS) burst criteria adapted from our previous study (Claar et al. 2023): burst onset required a preceding quiescent period of at least 100 ms (pre-burst ISI > 100 ms) followed by a first intra-burst ISI shorter than 5 ms, with subsequent spikes included if they occurred within 4 ms of the preceding spike (Grenier, Timofeev, and Steriade 1998; Contreras and Steriade 1995; Guido and Weyand 1995; Halassa et al. 2011; Lu, Guido, and Sherman 1992; Nestvogel and McCormick 2022). For cortical and other non-thalamic units, bursts were defined as three or more consecutive spikes with inter-spike intervals shorter than 10 ms, corresponding to the critical frequency for backpropagation-activated Ca²⁺ spike firing in layer 5 pyramidal neurons (Larkum, Zhu, and Sakmann 1999; de Kock and Sakmann 2008). To ensure each detected burst was a discrete event, we required a minimum preceding silence of 50 ms, following the MaxInterval framework (Cotterill and Eglen 2019; Bakkum et al. 2013) and consistent with the view that silence is a functionally distinct state demarcating burst onset (Friedenberger et al. 2023).

Four metrics were computed for each unit within each 5-minute time bin: *proportion of bursting units* (number of units that exhibit at least a burst over all the units for each selected area), *burst rate* (number of bursts per second of analyzed rest time), *mean burst size* (average number of spikes per burst), and *burst fraction*, defined as the proportion of spikes belonging to a burst out of the total number of spikes (N_spikes_in_bursts / N_total_spikes), in line with the definition of Oswald et al. (2007) (Oswald, Doiron, and Maler 2007). Metrics were computed in non-overlapping 5-minute bins spanning the last 10 min before the first injection as baseline and the post-second-injection period (up to 60 min post-injection). For each unit, the mean value of each burst metric across all available baseline bins served as the baseline reference, and post-injection values were expressed as absolute change from baseline:

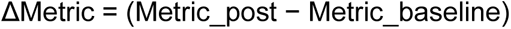

For burst rate, burst fraction, and mean burst size, time bins in which a unit had rest time but no detectable bursts were excluded from that bin’s group mean, so that these three metrics reflect burst intensity among actively bursting units; changes in the fraction of units that burst at all - including units newly recruited into bursting by the drug - were instead captured by the proportion of bursting units metric, which included all units with recorded rest time regardless of bursting status. Analyses were conducted at the level of major brain regions, cortical layers, and individual thalamic nuclei, as assigned from CCFv3 structure annotations. As in the firing rate analysis described below, epochs within a −100 to +600 ms window around each electrical or visual stimulation onset were excluded prior to burst detection, and the same inclusion criteria were applied: 1) a minimum of three subjects per condition, and 2) a minimum of 10 units from the brain area of interest, yielding a minimum of 10 units per brain area from at least 3 different subjects per condition. To summarize the drug effect over the period of peak action, we also computed a time-averaged contrast by collapsing across the 5-minute bins spanning 40-60 min post-injection. The resulting distributions of drug and saline conditions were compared between conditions and are displayed as violin plots. P-values for these comparisons were corrected for multiple comparisons using the Benjamini–Hochberg FDR procedure, as described in the Statistics section. Burst analysis was restricted to putative excitatory (RS) neurons, classified by spike waveform duration as described above, and performed exclusively during resting periods, defined as continuous epochs of at least 1 second with a locomotion speed below 1 cm/s. The RS/FS classification was not applied to striatal units or to units recorded in the reticular nucleus of the thalamus, where all isolated units were retained for analysis regardless of waveform duration.

#### Firing Rate Analysis

To quantify the effect of psilocybin on spontaneous spiking activity, we computed the mean firing rate of each unit across brain regions and experimental conditions. Firing rate was calculated as the total number of spikes divided by the total duration of analyzed rest time within each non-overlapping 5-minute bin. Analysis was performed exclusively during resting periods, defined as continuous epochs of at least 1 second with a locomotion speed below 1 cm/s. Epochs within a −100 to +600 ms window around each electrical or visual stimulation onset were excluded prior to firing rate computation.

Bins span the last 10 min before the first injection as baseline and the post-second-injection period (up to 60 min post-injection), yielding two baseline bins and up to twelve post-injection bins per session. For each unit, the mean firing rate across all available baseline bins served as the baseline reference, and post-injection values were expressed as the absolute change from baseline:

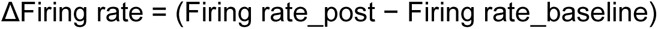

To isolate drug-specific effects, the drug–saline contrast was computed at each time bin and assessed for significance using the hierarchical bootstrap procedure described in the Statistics section. The same inclusion criteria were applied as for the burst analysis: a minimum of three subjects and ten units per brain area per condition. Units were included regardless of whether they were active during all bins; bins with no rest time were assigned to a firing rate of zero and excluded from the baseline mean. Analyses were conducted at the level of major brain regions, cortical areas and layers, thalamic nuclei groups, and hippocampal subregions, as described in the Neuroanatomical classification and region groupings section.

Similar to the burst analysis to summarize the drug effect over the period of peak action, we also computed a time-averaged contrast by collapsing across the 5-minute bins spanning 40-60 min post-injection. The resulting distributions of drug and saline conditions were compared between conditions and are displayed as violin plots.

All firing rate analyses were conducted separately for regular-spiking (RS) and fast-spiking (FS) units, classified based on spike waveform duration as described above.

#### Correlational analysis

For each unit, the change in burst rate and firing rate was defined as the difference between the mean value across the 40-60 min post-injection window and the unit’s pre-injection baseline mean. Units were included only if both metrics yielded a computable value. The Pearson correlation coefficient between the two change metrics was computed separately for each region and condition, with 95% confidence intervals derived via Fisher z-transformation and significance assessed by the associated two-tailed p-value; each unit contributed one observation. P-values were corrected for multiple comparisons across regions within each condition using the Benjamini–Hochberg false discovery rate procedure.

#### EEG and LFP spectral power analysis

Spectral power analysis was conducted during spontaneous activity periods, defined as intervals free of electrical stimulation lasting at least 10 consecutive s and not overlapping with a 60-second window before or after any injection. EEG and LFP signals recorded during these periods were segmented into non-overlapping 10-second epochs. For each subject and condition (saline, psilocybin, or Ket+Psi), data were included in further analysis only if a minimum of 9 valid 10-second epochs were present in both the baseline period (defined as the period prior to the first injection, common to all conditions) and the post-second-injection period. Subjects not meeting this criterion were excluded from spectral analysis (LFP final numbers: 45 sessions from 34 mice; EEG final numbers: 39 sessions from 29 mice).

For all remaining recordings, each 10-second epoch was bandpass filtered using a 3rd-order Butterworth filter (scipy.signal.butter and scipy.signal.filtfilt, SciPy, Python) across the following frequency bands: delta (0.1–4 Hz), theta (4–8 Hz), alpha (8–13 Hz), beta (13–35 Hz), gamma (35–55 Hz), and high gamma (65–100 Hz). Instantaneous amplitude was estimated by computing the Hilbert transform of each bandpass-filtered signal, and the absolute amplitude was averaged across each epoch to yield a measure of absolute power per frequency band. For EEG data, power was computed separately for each electrode channel. For LFP data, absolute Hilbert amplitudes were averaged across channels within each cortical layer and brain area prior to further analysis.

Median power was then computed across epochs within two-time windows: the baseline period (prior to the first injection) and the post-drug period (20–60 min after the second injection). The median percentage change in power relative to baseline for each frequency band was calculated as:

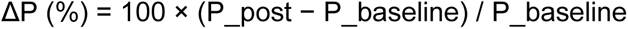

where P_post is the median power after the second injection during the saline, psilocybin, or Ket+Psi period, and P_baseline is the median power during baseline period. To compare spectral power changes between drug and saline conditions, the median percentage change was computed across subjects for each condition, and the difference between the drug condition (psilocybin or Ket+Psi) and the saline condition was calculated.

When behavioral state was taken into account, epochs were classified as resting if the mean locomotion speed over the full 10-second epoch was less than 1 cm/s, or as active if the mean speed exceeded 1 cm/s. Locomotion speed was derived from the running wheel’s angular velocity.

### Statistics

#### Behavioral effects

For population-level comparisons across all sessions, mean locomotion speed and mean normalized pupil radius during the post-injection period were compared to baseline within each condition using a Wilcoxon signed-rank test, and differences in these metrics between conditions (saline vs. psilocybin and saline vs. Ket+Psi) were assessed using a Wilcoxon rank-sum test. Multiple comparisons were corrected using Bonferroni correction.

For paired sessions, a linear mixed-effects model (LMM) was fit separately for the saline–psilocybin pairs and the saline–Ket+Psi pairs to assess the effect of drug type, time point (baseline/post-drug), their interaction, locomotion speed (z-scored), and time in experiment (z-scored) on continuous normalized pupil radius, with mouse identity included as a random effect. Significance of fixed-effect coefficients was assessed using Satterthwaite’s method for degrees of freedom approximation.

#### Spectral power statistical analysis

Spectral power in each frequency band during the post-injection period was compared to baseline using a Wilcoxon signed-rank test (pingouin.wilcoxon, Python) separately for each condition. When the multiple comparisons were considered, results were corrected for multiple comparisons across EEG channels (N=30) and across cortical areas and layers for LFP (N=11) using the Benjamini-Hochberg procedure (pingouin.multicomp, Python). To compare percentage power changes between conditions (saline vs. psilocybin or saline vs. Ket+Psi), a Mann-Whitney U test (pingouin.mwu, Python) was used, with the same multiple comparison corrections applied.

#### Units’ statistics

To assess whether psilocybin or Ket+Psi significantly altered firing rates and burst dynamics relative to saline at the population level and within individual brain regions, we used a hierarchical bootstrap procedure (Chen et al. 2024; Saravanan, Berman, and Sober 2020). Briefly, for each comparison, we resampled sessions with replacement at the first level and units with replacement at the second level, computing the mean difference in the metric of interest across B = 10,000 bootstrap iterations. A two-tailed p-value was derived from the bootstrap distribution as the proportion of resampled mean differences exceeding zero in the direction opposite to the observed effect, multiplied by two. This hierarchical approach accounts for the non-independence of units recorded within the same session. An exception applies to the proportion of bursting units metric, which is computed as a session-level aggregate: the fraction of all recorded units in a given region that exhibit at least one burst within a time bin. Because this metric is already averaged across units within each session, the bootstrap was applied at the session level only, resampling sessions with replacement without a second unit-level resampling step. The same session-level approach was used for all statistical comparisons involving this metric. Comparisons were performed at the level of major brain regions (isocortex, hippocampal formation, olfactory areas, striatum, and thalamus) as well as anatomical subregions and cortical layers as defined above. For heatmaps displaying the drug–saline contrast across 5-minute time bins spanning 0–60 min after the second injection, p-values were not corrected for multiple comparisons, consistent with the exploratory nature of the region-specific time-resolved analyses. For the distribution plots displaying the drug–saline contrast averaged across the 40-60 min post-injection window, p-values were corrected for multiple comparisons using the Benjamini–Hochberg false discovery rate (FDR) procedure.

## CODE AND DATA AVAILABILITY

Python scripts generated for the units’ analyses are available in the Github repository at the following link: https://github.com/Davi1990/psilocybin-burst-firing.

The raw data analyzed for this manuscript will be made publicly available in Neurodata Without Borders (NWB) format in a DANDI Archive upon acceptance for publication.

## ACKNOWLEDGMENTS

We wish to thank Jerome Lecoq and Andrew Shelton for insightful feedback, comments and useful discussions. We thank Yiliu Wang for feedback and inputs on the analytical approach. We thank the Allen Institute Animal Care, the Neurosurgery and Behavior, and the Lab Animal Services teams for mouse husbandry, care, and habituation; the Allen Institute Manufacturing and Process Engineering team for experimental hardware and software support. We also wish to thank the PostBac program of the Allen Institute for supporting Y.N.’s internship. We wish to thank the Allen Institute founder, Paul G. Allen, for his vision, encouragement, and support.

We gratefully acknowledge funding from the Templeton World Charity Foundation (TWCF-2022-30262; I.R.), the Tiny Blue Dot Foundation (Santa Monica, California, US), and the Allen Institute. DAP was partially supported by P50 MH119467.

## AUTHOR CONTRIBUTIONS STATEMENT

Conceptualization: I.R.; Methodology: I.R., C.K., L.C.; Software: L.C., I.R.; Validation: I.R., D.M., D.W., R.D.F.; Formal Analysis: D.M., Y.N., D.W., R.D.F., I.R.; Investigation: L.M., I.R., L.C.; Resources: C.K., I.R., T.O., D.A.P., M.B.; Data Curation: L.M., P.S., Y.N., L.C., D.W., I.R.; Writing – Original Draft: I.R., D.M., Y.N., L.M., L.C., D.W., R.D.F.; Writing – Review & Editing: I.R., C.K., T.O., D.A.P., L.M., Y.N., D.M., D.W., R.D.F., L.C., P.S., M.B.; Visualization: D.M., Y.N., D.W., L.C., R.D.F., I.R.; Supervision: I.R., C.K., T.O., M.B., D.A.P.; Project administration: I.R., L.M.; Funding Acquisition: I.R.

## REFERENCES

Alkire, M. T., A. G. Hudetz, and G. Tononi. 2008. ’Consciousness and anesthesia’, *Science*, 322: 876-80. Arnsten, A. F. 2009. ’Stress signalling pathways that impair prefrontal cortex structure and function’, Nat Rev Neurosci, 10: 410–22.

Alkire, M. T., A. G. Hudetz, and G. Tononi. 2015. ’Stress weakens prefrontal networks: molecular insults to higher cognition’, Nat Neurosci, 18: 1376–85.

Bains, J. S., J. M. Longacher, and K. J. Staley. 1999. ’Reciprocal interactions between CA3 network activity and strength of recurrent collateral synapses’, Nat Neurosci, 2: 720–6.

Bair, W., C. Koch, W. Newsome, and K. Britten. 1994. ’Power spectrum analysis of bursting cells in area MT in the behaving monkey’, J Neurosci, 14: 2870–92.

Bakkum, D. J., M. Radivojevic, U. Frey, F. Franke, A. Hierlemann, and H. Takahashi. 2013. ’Parameters for burst detection’, Front Comput Neurosci, 7: 193.

Barrientos, R., A. Alatorre, A. Oviedo-Chavez, A. Delgado, N. Nielsen, and E. Querejeta. 2022. ’Tonic serotonergic input increases the burst firing mode and diminishes the firing rate of reticular thalamic nucleus neurons through 5-HT1A receptors activation in anesthetized rats’, Exp Brain Res, 240: 1341–56.

Bartho, P., H. Hirase, L. Monconduit, M. Zugaro, K. D. Harris, and G. Buzsaki. 2004. ’Characterization of neocortical principal cells and interneurons by network interactions and extracellular features’, J Neurophysiol, 92: 600–8.

Bartho, P., A. Slezia, F. Matyas, L. Faradzs-Zade, I. Ulbert, K. D. Harris, and L. Acsady. 2014. ’Ongoing network state controls the length of sleep spindles via inhibitory activity’, Neuron, 82: 1367–79.

Bortone, D. S., S. R. Olsen, and M. Scanziani. 2014. ’Translaminar inhibitory cells recruited by layer 6 corticothalamic neurons suppress visual cortex’, Neuron, 82: 474–85.

Breant, B. J. B., J. Prius Mengual, A. Andrews, A. Hoerder-Suabedissen, J. Patel, D. M. Bannerman, T. Sharp, and V. V. Vyazovskiy. 2026. ’Vigilance state dissociation induced by 5-MeO-DMT in mice’, Commun Biol, 9: 163.

Bruno, R. M., and D. J. Simons. 2002. ’Feedforward mechanisms of excitatory and inhibitory cortical receptive fields’, J Neurosci, 22: 10966–75.

Carbonaro, T. M., M. W. Johnson, E. Hurwitz, and R. R. Griffiths. 2018. ’Double-blind comparison of the two hallucinogens psilocybin and dextromethorphan: similarities and differences in subjective experiences’, Psychopharmacology (Berl), 235: 521–34.

Carhart-Harris, R. L., D. Erritzoe, T. Williams, J. M. Stone, L. J. Reed, A. Colasanti, R. J. Tyacke, R. Leech, A. L. Malizia, K. Murphy, P. Hobden, J. Evans, A. Feilding, R. G. Wise, and D. J. Nutt. 2012. ’Neural correlates of the psychedelic state as determined by fMRI studies with psilocybin’, Proc Natl Acad Sci U S A, 109: 2138–43.

Carhart-Harris, R. L., and K. J. Friston. 2019. ’REBUS and the Anarchic Brain: Toward a Unified Model of the Brain Action of Psychedelics’, Pharmacol Rev, 71: 316–44.

Carhart-Harris, R. L., L. Roseman, M. Bolstridge, L. Demetriou, J. N. Pannekoek, M. B. Wall, M. Tanner, M. Kaelen, J. McGonigle, K. Murphy, R. Leech, H. V. Curran, and D. J. Nutt. 2017. ’Psilocybin for treatment-resistant depression: fMRI-measured brain mechanisms’, Sci Rep, 7: 13187.

Catlow, B. J., S. Song, D. A. Paredes, C. L. Kirstein, and J. Sanchez-Ramos. 2013. ’Effects of psilocybin on hippocampal neurogenesis and extinction of trace fear conditioning’, Exp Brain Res, 228: 481–91.

Chen, S., Y. Liu, Z. A. Wang, J. Colonell, L. D. Liu, H. Hou, N. W. Tien, T. Wang, T. Harris, S. Druckmann, N. Li, and K. Svoboda. 2024. ’Brain-wide neural activity underlying memory-guided movement’, Cell, 187: 676–91 e16.

Claar, L. D., I. Rembado, J. R. Kuyat, S. Russo, L. C. Marks, S. R. Olsen, and C. Koch. 2023. ’Cortico-thalamo-cortical interactions modulate electrically evoked EEG responses in mice’, Elife, 12.

Contreras, D., and M. Steriade. 1995. ’Cellular basis of EEG slow rhythms: a study of dynamic corticothalamic relationships’, J Neurosci, 15: 604–22.

Corlett, P. R., G. Horga, P. C. Fletcher, B. Alderson-Day, K. Schmack, and A. R. Powers, 3rd. 2019. ’Hallucinations and Strong Priors’, Trends Cogn Sci, 23: 114–27.

Cotterill, E., and S. J. Eglen. 2019. ’Burst Detection Methods’, Adv Neurobiol, 22: 185–206.

Crick, F. 1984. ’Function of the thalamic reticular complex: the searchlight hypothesis’, Proc Natl Acad Sci U S A, 81: 4586–90.

Daws, R. E., C. Timmermann, B. Giribaldi, J. D. Sexton, M. B. Wall, D. Erritzoe, L. Roseman, D. Nutt, and R. Carhart-Harris. 2022. ’Increased global integration in the brain after psilocybin therapy for depression’, Nat Med, 28: 844–51.

De Filippo, R., R. Gillis, D. Wyrick, M. Carlson, S. Durand, R.C. Peene, A. Bawany, A. Amaya, H. Belski, C. Grasso, W. Han, J. Kenney, C. Kiselycznyk, H. Loeffler, L. Marks, R. Naidoo, B. Ouellette, L. Suarez, J. Swapp, T. Johnson, J. Weber, J. Wilkes, P.A. Groblewski, A. Williford, M. Buice, C. Koch, I. Rembado, J. Lecoq, and T. Ott. Under submission. ’Psilocybin collapses visual change detection and drives cortical dynamics toward a state of surprise’.

de Kock, C. P., and B. Sakmann. 2008. ’High frequency action potential bursts (>or= 100 Hz) in L2/3 and L5B thick tufted neurons in anaesthetized and awake rat primary somatosensory cortex’, J Physiol, 586: 3353–64.

Doss, M. K., M. B. Madden, A. Gaddis, M. B. Nebel, R. R. Griffiths, B. N. Mathur, and F. S. Barrett. 2022. ’Models of psychedelic drug action: modulation of cortical-subcortical circuits’, Brain, 145: 441–56.

Ekins, T. G., I. Brooks, S. Kailasa, C. Rybicki-Kler, I. Jedrasiak-Cape, E. Donoho, G. A. Mashour, J. Rech, and O. J. Ahmed. 2023. ’Cellular rules underlying psychedelic control of prefrontal pyramidal neurons’, bioRxiv.

Fadahunsi, N., J. Lund, A. W. Breum, C. V. Mathiesen, I. B. Larsen, G. M. Knudsen, A. B. Klein, and C. Clemmensen. 2022. ’Acute and long-term effects of psilocybin on energy balance and feeding behavior in mice’, Transl Psychiatry, 12: 330.

Ferrarelli, F., and G. Tononi. 2011. ’The thalamic reticular nucleus and schizophrenia’, Schizophr Bull, 37: 306–15.

Friedenberger, Z., E. Harkin, K. Toth, and R. Naud. 2023. ’Silences, spikes and bursts: Three-part knot of the neural code’, J Physiol, 601: 5165–93.

Gaddis, A., D. E. Lidstone, M. B. Nebel, R. R. Griffiths, S. H. Mostofsky, A. F. Mejia, and F. S. Barrett. 2022. ’Psilocybin induces spatially constrained alterations in thalamic functional organizaton and connectivity’, Neuroimage, 260: 119434.

Geyer, M. A., and F. X. Vollenweider. 2008. ’Serotonin research: contributions to understanding psychoses’, Trends Pharmacol Sci, 29: 445–53.

Golden, C. T., and P. Chadderton. 2022. ’Psilocybin reduces low frequency oscillatory power and neuronal phase-locking in the anterior cingulate cortex of awake rodents’, Sci Rep, 12: 12702.

Gonzalo-Ruiz, A., A. R. Lieberman, and J. M. Sanz-Anquela. 1995. ’Organization of serotoninergic projections from the raphe nuclei to the anterior thalamic nuclei in the rat: a combined retrograde tracing and 5-HT immunohistochemical study’, J Chem Neuroanat, 8: 103–15.

Grenier, F., I. Timofeev, and M. Steriade. 1998. ’Leading role of thalamic over cortical neurons during postinhibitory rebound excitation’, Proc Natl Acad Sci U S A, 95: 13929–34.

Groblewski, P. A., D. Sullivan, J. Lecoq, S. E. J. de Vries, S. Caldejon, Q. L’Heureux, T. Keenan, K. Roll, C. Slaughterback, A. Williford, and C. Farrell. 2020. ’A standardized head-fixation system for performing large-scale, in vivo physiological recordings in mice’, J Neurosci Methods, 346: 108922.

Guido, W., and T. Weyand. 1995. ’Burst responses in thalamic relay cells of the awake behaving cat’, J Neurophysiol, 74: 1782–6.

Guo, F., Q. Zhang, B. Zhang, Z. Fu, B. Wu, C. Huang, and Y. Li. 2014. ’Burst-firing patterns in the prefrontal cortex underlying the neuronal mechanisms of depression probed by antidepressants’, Eur J Neurosci, 40: 3538–47.

Guo, Z. V., H. K. Inagaki, K. Daie, S. Druckmann, C. R. Gerfen, and K. Svoboda. 2017. ’Maintenance of persistent activity in a frontal thalamocortical loop’, Nature, 545: 181–86.

Halassa, M. M., and S. Kastner. 2017. ’Thalamic functions in distributed cognitive control’, Nat Neurosci, 20: 1669–79.

Halassa, M. M., J. H. Siegle, J. T. Ritt, J. T. Ting, G. Feng, and C. I. Moore. 2011. ’Selective optical drive of thalamic reticular nucleus generates thalamic bursts and cortical spindles’, Nat Neurosci, 14: 1118–20.

Halberstadt, A. L. 2015. ’Recent advances in the neuropsychopharmacology of serotonergic hallucinogens’, Behav Brain Res, 277: 99–120.

Harris, J. A., S. Mihalas, K. E. Hirokawa, J. D. Whitesell, H. Choi, A. Bernard, P. Bohn, S. Caldejon, L. Casal, A. Cho, A. Feiner, D. Feng, N. Gaudreault, C. R. Gerfen, N. Graddis, P. A. Groblewski, A. M. Henry, A. Ho, R. Howard, J. E. Knox, L. Kuan, X. Kuang, J. Lecoq, P. Lesnar, Y. Li, J. Luviano, S. McConoughey, M. T. Mortrud, M. Naeemi, L. Ng, S. W. Oh, B. Ouellette, E. Shen, S. A. Sorensen, W. Wakeman, Q. Wang, Y. Wang, A. Williford, J. W. Phillips, A. R. Jones, C. Koch, and H. Zeng. 2019. ’Hierarchical organization of cortical and thalamic connectivity’, Nature, 575: 195–202.

Harris, K. D., H. Hirase, X. Leinekugel, D. A. Henze, and G. Buzsaki. 2001. ’Temporal interaction between single spikes and complex spike bursts in hippocampal pyramidal cells’, Neuron, 32: 141–9.

Hesselgrave, N., T. A. Troppoli, A. B. Wulff, A. B. Cole, and S. M. Thompson. 2021. ’Harnessing psilocybin: antidepressant-like behavioral and synaptic actions of psilocybin are independent of 5-HT2R activation in mice’, Proc Natl Acad Sci U S A, 118.

Hidalgo Jimenez, J., K. Kristjan Kaup, and J. Aru. 2026. ’Electrophysiological mechanisms of psychedelic drugs: A systematic review’, Neurosci Biobehav Rev, 185: 106649.

Hintiryan, H., M. Rudd, S. Nanda, A. E. Gutierrez, D. Lo, T. Boesen, L. Garcia, J. Sun, C. Estrada, H. S. Mun, S. Yamashita, Y. E. Han, I. Bowman, L. Gou, C. Cao, J. Gonzalez, K. Moradi, Q. Zhao, I. Yenokian, A. Dev, B. Zingg, H. Xu, Q. Xue, M. Zhu, L. Liu, X. Chen, Z. Yun, H. Peng, N. N. Foster, and H. W. Dong. 2025. ’Distinct subnetworks of the mouse anterior thalamic nuclei’, Nat Commun, 16: 6018.

Huo, Y., H. Chen, and Z. V. Guo. 2020. ’Mapping Functional Connectivity from the Dorsal Cortex to the Thalamus’, Neuron, 107: 1080–94 e5.

Jiang, Q., L. X. Shao, S. Yao, N. K. Savalia, A. D. Gilbert, P. A. Davoudian, J. D. Nothnagel, G. Tian, T. S. Hung, H. M. Lai, K. T. Beier, H. Zeng, and A. C. Kwan. 2026. ’Psilocybin triggers an activity-dependent rewiring of large-scale cortical networks’, Cell, 189: 659–75 e22.

Juczewski, K., J. A. Koussa, A. J. Kesner, J. O. Lee, and D. M. Lovinger. 2020. ’Stress and behavioral correlates in the head-fixed method: stress measurements, habituation dynamics, locomotion, and motor-skill learning in mice’, Sci Rep, 10: 12245.

Jun, J. J., N. A. Steinmetz, J. H. Siegle, D. J. Denman, M. Bauza, B. Barbarits, A. K. Lee, C. A. Anastassiou, A. Andrei, C. Aydin, M. Barbic, T. J. Blanche, V. Bonin, J. Couto, B. Dutta, S. L. Gratiy, D. A. Gutnisky, M. Hausser, B. Karsh, P. Ledochowitsch, C. M. Lopez, C. Mitelut, S. Musa, M. Okun, M. Pachitariu, J. Putzeys, P. D. Rich, C. Rossant, W. L. Sun, K. Svoboda, M. Carandini, K. D. Harris, C. Koch, J. O’Keefe, and T. D. Harris. 2017. ’Fully integrated silicon probes for high-density recording of neural activity’, Nature, 551: 232–36.

Kasper, E. M., A. U. Larkman, J. Lubke, and C. Blakemore. 1994. ’Pyramidal neurons in layer 5 of the rat visual cortex. I. Correlation among cell morphology, intrinsic electrophysiological properties, and axon targets’, J Comp Neurol, 339: 459–74.

Kheirbek, M. A., K. C. Klemenhagen, A. Sahay, and R. Hen. 2012. ’Neurogenesis and generalization: a new approach to stratify and treat anxiety disorders’, Nat Neurosci, 15: 1613–20.

Koch, Christof. 1999. *Biophysics of computation : information processing in single neurons* (Oxford University Press: New York).

Kometer, M., A. Schmidt, L. Jancke, and F. X. Vollenweider. 2013. ’Activation of serotonin 2A receptors underlies the psilocybin-induced effects on alpha oscillations, N170 visual-evoked potentials, and visual hallucinations’, J Neurosci, 33: 10544–51.

Kumar, V. J., K. Scheffler, and W. Grodd. 2023. ’The structural connectivity mapping of the intralaminar thalamic nuclei’, Sci Rep, 13: 11938.

Kwan, A. C., D. E. Olson, K. H. Preller, and B. L. Roth. 2022. ’The neural basis of psychedelic action’, Nat Neurosci, 25: 1407–19.

Larkum, M. 2013. ’A cellular mechanism for cortical associations: an organizing principle for the cerebral cortex’, Trends Neurosci, 36: 141–51.

Larkum, M. E., J. J. Zhu, and B. Sakmann. 1999. ’A new cellular mechanism for coupling inputs arriving at different cortical layers’, Nature, 398: 338–41.

Leow, Y. N., B. Zhou, H. A. Sullivan, A. R. Barlowe, I. R. Wickersham, and M. Sur. 2022. ’Brain-wide mapping of inputs to the mouse lateral posterior (LP/Pulvinar) thalamus-anterior cingulate cortex network’, J Comp Neurol, 530: 1992–2013.

Lisman, J. E. 1997. ’Bursts as a unit of neural information: making unreliable synapses reliable’, Trends Neurosci, 20: 38–43.

Liu, L. D., S. Chen, H. Hou, S. J. West, M. Faulkner, Laboratory International Brain, M. N. Economo, N. Li, and K. Svoboda. 2021. ’Accurate Localization of Linear Probe Electrode Arrays across Multiple Brains’, eNeuro, 8.

Llinas, R. R., U. Ribary, D. Jeanmonod, E. Kronberg, and P. P. Mitra. 1999. ’Thalamocortical dysrhythmia: A neurological and neuropsychiatric syndrome characterized by magnetoencephalography’, Proc Natl Acad Sci U S A, 96: 15222–7.

Llinas, R., U. Ribary, D. Contreras, and C. Pedroarena. 1998. ’The neuronal basis for consciousness’, Philos Trans R Soc Lond B Biol Sci, 353: 1841–9.

Logothetis, N. K. 2008. ’What we can do and what we cannot do with fMRI’, *Nature*, 453: 869-78. Lopez-Gimenez, J. F., and J. Gonzalez-Maeso. 2018. ’Hallucinogens and Serotonin 5-HT(2A) Receptor-Mediated Signaling Pathways’, Curr Top Behav Neurosci, 36: 45–73.

Lu, O. D., K. White, K. Raymond, C. Liu, A. S. Klein, N. Green, S. Vaillancourt, A. Gallagher, L. Shindy, A. Li, K. Wallquist, R. Li, M. Zou, A. B. Casey, L. P. Cameron, M. B. Pomrenze, V. Sohal, M. A. Kheirbek, A. M. Gomez, S. Lammel, B. D. Heifets, and R. Malenka. 2025. ’A multi-institutional investigation of psilocybin’s effects on mouse behavior’, bioRxiv.

Lu, S. M., W. Guido, and S. M. Sherman. 1992. ’Effects of membrane voltage on receptive field properties of lateral geniculate neurons in the cat: contributions of the low-threshold Ca2+ conductance’, J Neurophysiol, 68: 2185–98.

Ly, C., A. C. Greb, L. P. Cameron, J. M. Wong, E. V. Barragan, P. C. Wilson, K. F. Burbach, S. Soltanzadeh Zarandi, A. Sood, M. R. Paddy, W. C. Duim, M. Y. Dennis, A. K. McAllister, K. M. Ori-McKenney, J. A. Gray, and D. E. Olson. 2018. ’Psychedelics Promote Structural and Functional Neural Plasticity’, Cell Rep, 23: 3170–82.

Magee, J. C., and D. Johnston. 1997. ’A synaptically controlled, associative signal for Hebbian plasticity in hippocampal neurons’, Science, 275: 209–13.

Marks, L. C.;, I.; Rembado, and L. D Claar. ’Simultaneous Recording of Neuropixels and EEG in Head-Fixed Mice.’, protocols.io 10.17504/protocols.io.14egn9q16l5d/v1.

Marlinski, V., and I. N. Beloozerova. 2014. ’Burst firing of neurons in the thalamic reticular nucleus during locomotion’, J Neurophysiol, 112: 181–92.

Mathis, A., P. Mamidanna, K. M. Cury, T. Abe, V. N. Murthy, M. W. Mathis, and M. Bethge. 2018. ’DeepLabCut: markerless pose estimation of user-defined body parts with deep learning’, Nat Neurosci, 21: 1281–89.

Muller, F., C. Lenz, P. Dolder, U. Lang, A. Schmidt, M. Liechti, and S. Borgwardt. 2017. ’Increased thalamic resting-state connectivity as a core driver of LSD-induced hallucinations’, Acta Psychiatr Scand, 136: 648–57.

Nakazawa, K., M. C. Quirk, R. A. Chitwood, M. Watanabe, M. F. Yeckel, L. D. Sun, A. Kato, C. A. Carr, D. Johnston, M. A. Wilson, and S. Tonegawa. 2002. ’Requirement for hippocampal CA3 NMDA receptors in associative memory recall’, Science, 297: 211–8.

Nelson, A. J. D. 2021. ’The anterior thalamic nuclei and cognition: A role beyond space?’, Neurosci Biobehav Rev, 126: 1–11.

Nestvogel, D. B., and D. A. McCormick. 2022. ’Visual thalamocortical mechanisms of waking state-dependent activity and alpha oscillations’, Neuron, 110: 120–38 e4.

Niell, C. M., and M. P. Stryker. 2008. ’Highly selective receptive fields in mouse visual cortex’, J Neurosci, 28: 7520–36.

Nutt, D., D. Erritzoe, and R. Carhart-Harris. 2020. ’Psychedelic Psychiatry’s Brave New World’, Cell, 181: 24–28.

Oh, S. W., J. A. Harris, L. Ng, B. Winslow, N. Cain, S. Mihalas, Q. Wang, C. Lau, L. Kuan, A. M. Henry, M. T. Mortrud, B. Ouellette, T. N. Nguyen, S. A. Sorensen, C. R. Slaughterbeck, W. Wakeman, Y. Li, D. Feng, A. Ho, E. Nicholas, K. E. Hirokawa, P. Bohn, K. M. Joines, H. Peng, M. J. Hawrylycz, J. W. Phillips, J. G. Hohmann, P. Wohnoutka, C. R. Gerfen, C. Koch, A. Bernard, C. Dang, A. R. Jones, and H. Zeng. 2014. ’A mesoscale connectome of the mouse brain’, Nature, 508: 207–14.

Onofrj, M., M. Russo, S. Delli Pizzi, D. De Gregorio, A. Inserra, G. Gobbi, and S. L. Sensi. 2023. ’The central role of the Thalamus in psychosis, lessons from neurodegenerative diseases and psychedelics’, Transl Psychiatry, 13: 384.

Oswald, A. M., B. Doiron, and L. Maler. 2007. ’Interval coding. I. Burst interspike intervals as indicators of stimulus intensity’, J Neurophysiol, 97: 2731–43.

Passie, T., J. Seifert, U. Schneider, and H. M. Emrich. 2002. ’The pharmacology of psilocybin’, Addict Biol, 7: 357–64.

Pike, F. G., R. M. Meredith, A. W. Olding, and O. Paulsen. 1999. ’Rapid report: postsynaptic bursting is essential for ’Hebbian’ induction of associative long-term potentiation at excitatory synapses in rat hippocampus’, J Physiol, 518 (Pt 2): 571–6.

Pines, A. R., X. Zhang, J. Kochalka, S. S. Vesuna, I. V. Kauvar, D. Rajasekharan, T. R. Reneau, T. J. Akiki, L. M. Hack, J. S. Siegel, and L. M. Williams. 2026. ’Psychedelics disrupt hierarchical cortical propagations in the default mode network of humans and mice’, Proc Natl Acad Sci U S A, 123: e2522000123.

Pompeiano, M., J. M. Palacios, and G. Mengod. 1994. ’Distribution of the serotonin 5-HT2 receptor family mRNAs: comparison between 5-HT2A and 5-HT2C receptors’, Brain Res Mol Brain Res, 23: 163–78.

Preller, K. H., J. B. Burt, J. L. Ji, C. H. Schleifer, B. D. Adkinson, P. Stampfli, E. Seifritz, G. Repovs, J. H. Krystal, J. D. Murray, F. X. Vollenweider, and A. Anticevic. 2018. ’Changes in global and thalamic brain connectivity in LSD-induced altered states of consciousness are attributable to the 5-HT2A receptor’, Elife, 7.

Preller, K. H., A. Razi, P. Zeidman, P. Stampfli, K. J. Friston, and F. X. Vollenweider. 2019. ’Effective connectivity changes in LSD-induced altered states of consciousness in humans’, Proc Natl Acad Sci U S A, 116: 2743–48.

Purple, R. J., R. Gupta, C. W. Thomas, C. T. Golden, N. Palomero-Gallagher, R. Carhart-Harris, S. Froudist-Walsh, and M. W. Jones. 2025. ’Short- and long-term modulation of rat prefrontal cortical activity following single doses of psilocybin’, Mol Psychiatry, 30: 5889–900.

Ragan, T., L. R. Kadiri, K. U. Venkataraju, K. Bahlmann, J. Sutin, J. Taranda, I. Arganda-Carreras, Y. Kim, H. S. Seung, and P. Osten. 2012. ’Serial two-photon tomography for automated ex vivo mouse brain imaging’, Nat Methods, 9: 255–8.

Raison, C. L., G. Sanacora, J. Woolley, K. Heinzerling, B. W. Dunlop, R. T. Brown, R. Kakar, M. Hassman, R. P. Trivedi, R. Robison, N. Gukasyan, S. M. Nayak, X. Hu, K. C. O’Donnell, B. Kelmendi, J. Sloshower, A. D. Penn, E. Bradley, D. F. Kelly, T. Mletzko, C. R. Nicholas, P. R. Hutson, G. Tarpley, M. Utzinger, K. Lenoch, K. Warchol, T. Gapasin, M. C. Davis, C. Nelson-Douthit, S. Wilson, C. Brown, W. Linton, S. Ross, and R. R. Griffiths. 2023. ’Single-Dose Psilocybin Treatment for Major Depressive Disorder: A Randomized Clinical Trial’, JAMA, 330: 843–53.

Raus Balind, S., A. Mago, M. Ahmadi, N. Kis, Z. Varga-Nemeth, A. Lorincz, and J. K. Makara. 2019. ’Diverse synaptic and dendritic mechanisms of complex spike burst generation in hippocampal CA3 pyramidal cells’, Nat Commun, 10: 1859.

Raval, N. R., A. Johansen, L. L. Donovan, N. F. Ros, B. Ozenne, H. D. Hansen, and G. M. Knudsen. 2021. ’A Single Dose of Psilocybin Increases Synaptic Density and Decreases 5-HT(2A) Receptor Density in the Pig Brain’, Int J Mol Sci, 22.

Rodriguez, J. J., H. N. Noristani, W. B. Hoover, S. B. Linley, and R. P. Vertes. 2011. ’Serotonergic projections and serotonin receptor expression in the reticular nucleus of the thalamus in the rat’, Synapse, 65: 919–28.

Rolls, E. T. 2007. ’An attractor network in the hippocampus: theory and neurophysiology’, Learn Mem, 14: 714–31.

Roth, M. M., J. C. Dahmen, D. R. Muir, F. Imhof, F. J. Martini, and S. B. Hofer. 2016. ’Thalamic nuclei convey diverse contextual information to layer 1 of visual cortex’, Nat Neurosci, 19: 299–307.

Roy, R., and R. Narayanan. 2023. ’Ion-channel degeneracy and heterogeneities in the emergence of complex spike bursts in CA3 pyramidal neurons’, J Physiol, 601: 3297–328.

Russo, S., L. D. Claar, G. Furregoni, L. C. Marks, G. Krishnan, F. M. Zauli, G. Hassan, M. Solbiati, P. d’Orio, E. Mikulan, S. Sarasso, M. Rosanova, I. Sartori, M. Bazhenov, A. Pigorini, M. Massimini, C. Koch, and I. Rembado. 2025. ’Thalamic feedback shapes brain responses evoked by cortical stimulation in mice and humans’, Nat Commun, 16: 3627.

Saravanan, V., G. J. Berman, and S. J. Sober. 2020. ’Application of the hierarchical bootstrap to multi-level data in neuroscience’, Neuron Behav Data Anal Theory, 3.

Seeburg, D. P., X. Liu, and C. Chen. 2004. ’Frequency-dependent modulation of retinogeniculate transmission by serotonin’, J Neurosci, 24: 10950–62.

Seyfourian, P., L. C. Marks, L. D. Claar, Y. Nahas, M. Keating, C. Koch, and I. Rembado. 2026. ’Pupil-DLC: An open-source deep learning pipeline for scalable, marker-less tracking of pupil dynamics across conscious and unconscious states’, J Neurosci Methods, 435: 110848.

Shao, L. X., C. Liao, P. A. Davoudian, N. K. Savalia, Q. Jiang, C. Wojtasiewicz, D. Tan, J. D. Nothnagel, R. J. Liu, S. C. Woodburn, O. M. Bilash, H. Kim, A. Che, and A. C. Kwan. 2025. ’Psilocybin’s lasting action requires pyramidal cell types and 5-HT(2A) receptors’, Nature.

Shao, L. X., C. Liao, I. Gregg, P. A. Davoudian, N. K. Savalia, K. Delagarza, and A. C. Kwan. 2021. ’Psilocybin induces rapid and persistent growth of dendritic spines in frontal cortex in vivo’, Neuron, 109: 2535–44 e4.

Sherman, S. M. 2007. ’The thalamus is more than just a relay’, Curr Opin Neurobiol, 17: 417–22.

Sherman, S.M., and R. W. Guillery. 2006. ’Exploring the Thalamus and Its Role in Cortical Function’, MIT Press.

Siegle, J. H., X. Jia, S. Durand, S. Gale, C. Bennett, N. Graddis, G. Heller, T. K. Ramirez, H. Choi, J. A. Luviano, P. A. Groblewski, R. Ahmed, A. Arkhipov, A. Bernard, Y. N. Billeh, D. Brown, M. A. Buice, N. Cain, S. Caldejon, L. Casal, A. Cho, M. Chvilicek, T. C. Cox, K. Dai, D. J. Denman, S. E. J. de Vries, R. Dietzman, L. Esposito, C. Farrell, D. Feng, J. Galbraith, M. Garrett, E. C. Gelfand, N. Hancock, J. A. Harris, R. Howard, B. Hu, R. Hytnen, R. Iyer, E. Jessett, K. Johnson, I. Kato, J. Kiggins, S. Lambert, J. Lecoq, P. Ledochowitsch, J. H. Lee, A. Leon, Y. Li, E. Liang, F. Long, K. Mace, J. Melchior, D. Millman, T. Mollenkopf, C. Nayan, L. Ng, K. Ngo, T. Nguyen, P. R. Nicovich, K. North, G. K. Ocker, D. Ollerenshaw, M. Oliver, M. Pachitariu, J. Perkins, M. Reding, D. Reid, M. Robertson, K. Ronellenfitch, S. Seid, C. Slaughterbeck, M. Stoecklin, D. Sullivan, B. Sutton, J. Swapp, C. Thompson, K. Turner, W. Wakeman, J. D. Whitesell, D. Williams, A. Williford, R. Young, H. Zeng, S. Naylor, J. W. Phillips, R. C. Reid, S. Mihalas, S. R. Olsen, and C. Koch. 2021. ’Survey of spiking in the mouse visual system reveals functional hierarchy’, Nature, 592: 86–92.

Siegle, J. H., A. C. Lopez, Y. A. Patel, K. Abramov, S. Ohayon, and J. Voigts. 2017. ’Open Ephys: an open-source, plugin-based platform for multichannel electrophysiology’, J Neural Eng, 14: 045003.

Sirota, A., S. Montgomery, S. Fujisawa, Y. Isomura, M. Zugaro, and G. Buzsaki. 2008. ’Entrainment of neocortical neurons and gamma oscillations by the hippocampal theta rhythm’, Neuron, 60: 683–97.

Skyberg, R. J., C. W. Fields, D. M. Martins, and C. M. Niell. 2025. ’The impact of the serotonergic psychedelic DOI on active vision in freely moving mice’, bioRxiv.

Stringer, C., M. Pachitariu, N. Steinmetz, C. B. Reddy, M. Carandini, and K. D. Harris. 2019. ’Spontaneous behaviors drive multidimensional, brainwide activity’, Science, 364: 255.

Thomas, M. J., A. M. Watabe, T. D. Moody, M. Makhinson, and T. J. O’Dell. 1998. ’Postsynaptic complex spike bursting enables the induction of LTP by theta frequency synaptic stimulation’, J Neurosci, 18: 7118–26.

Van der Werf, Y. D., M. P. Witter, and H. J. Groenewegen. 2002. ’The intralaminar and midline nuclei of the thalamus. Anatomical and functional evidence for participation in processes of arousal and awareness’, Brain Res Brain Res Rev, 39: 107–40.

van Strien, N. M., N. L. Cappaert, and M. P. Witter. 2009. ’The anatomy of memory: an interactive overview of the parahippocampal-hippocampal network’, Nat Rev Neurosci, 10: 272–82.

Vertes, R. P., S. B. Linley, and W. B. Hoover. 2010. ’Pattern of distribution of serotonergic fibers to the thalamus of the rat’, Brain Struct Funct, 215: 1–28.

Vollenweider, F. X., and M. A. Geyer. 2001. ’A systems model of altered consciousness: integrating natural and drug-induced psychoses’, Brain Res Bull, 56: 495–507.

Vollenweider, F. X., and K. H. Preller. 2020. ’Psychedelic drugs: neurobiology and potential for treatment of psychiatric disorders’, Nat Rev Neurosci, 21: 611–24.

Vollenweider, F. X., M. F. Vollenweider-Scherpenhuyzen, A. Babler, H. Vogel, and D. Hell. 1998. ’Psilocybin induces schizophrenia-like psychosis in humans via a serotonin-2 agonist action’, Neuroreport, 9: 3897–902.

Wang, Z., and D. A. McCormick. 1993. ’Control of firing mode of corticotectal and corticopontine layer V burst-generating neurons by norepinephrine, acetylcholine, and 1S,3R-ACPD’, J Neurosci, 13: 2199–216.

Ward, L. M. 2011. ’The thalamic dynamic core theory of conscious experience’, Conscious Cogn, 20: 464–86.

