## Supplementary Material for "Brain-wide reconfiguration of burst firing by psilocybin reveals 5-HT2A-dependent circuit dynamics"

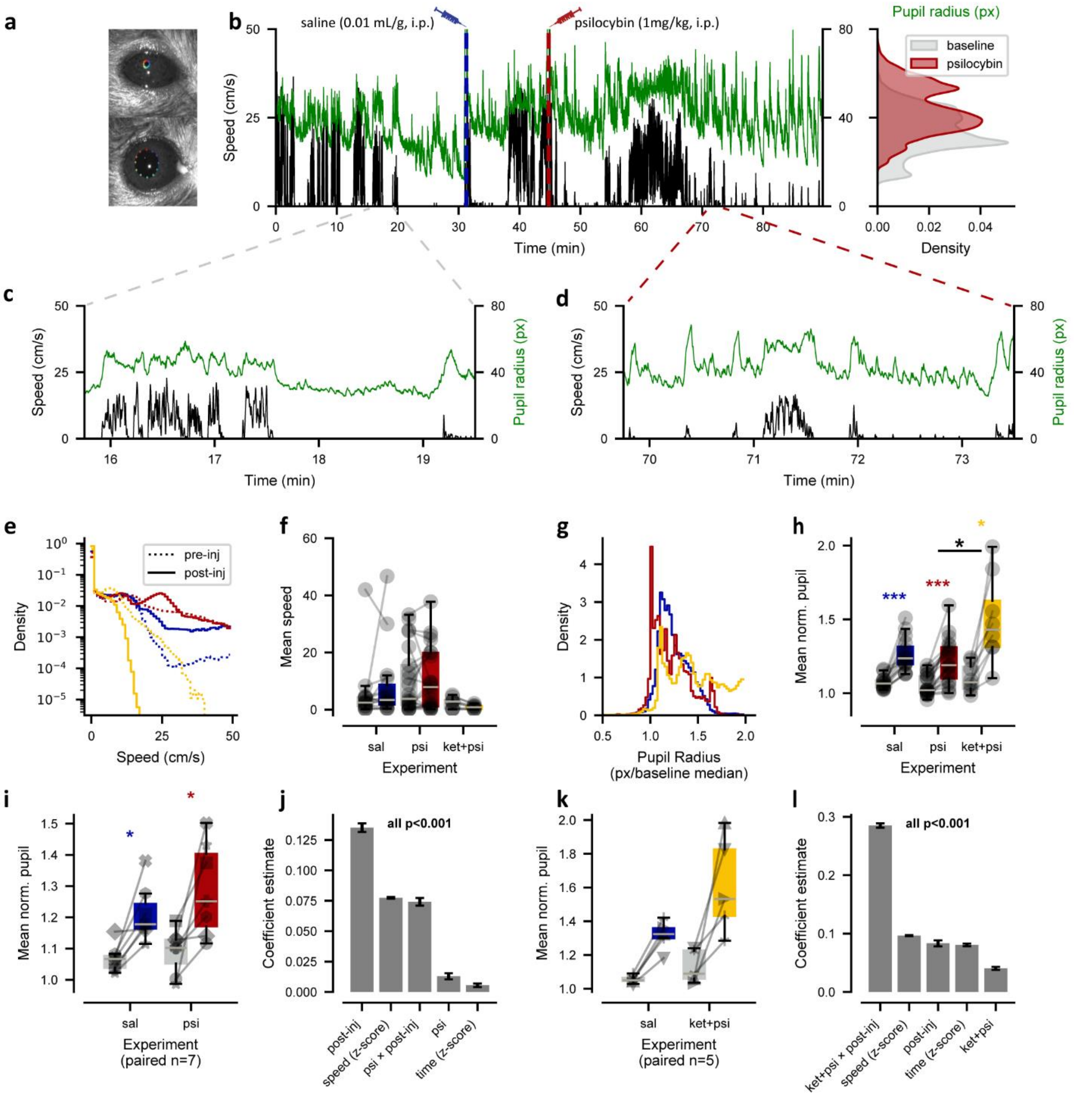


**Figure S1. Effects of psilocybin and Ket + Psi on locomotion and pupil size.**

a, Representative video frames showing automated pupil tracking using our tracking pipeline (Seyfourian et al, 2026), with 12 multicolor landmarks annotated around the pupil boundary. The model was trained to extract pupil radius across the full duration of each experimental session.

b, Running speed (black, left y-axis) and pupil radius (green, right y-axis) over time for one representative psilocybin experiment (same session as Figure 1e, trimmed to 90 min). Vertical dashed lines indicate the time of each injection (saline, blue; psilocybin, red). Right: distributions of pupil radius during the baseline period (prior to the first injection, gray) and the post-psilocybin period (after the second injection, red).

c, Expanded view of approximately 3 min immediately before the saline injection from b.

d, Expanded view of approximately 3 min following the psilocybin injection from b.

e, Running speed distributions for the 30–45 min post-second-injection window across saline (sal, n=15, blue), psilocybin (psi, n=21, red), and Ket + Psi (n=8, yellow) experiments. Dashed lines show the corresponding baseline speed distributions for the same sessions.

f, Mean running speed (30–45 min post-second injection) per session for each experimental condition; baseline means are shown in gray for all conditions.

g, Distributions of pupil radius normalized to the baseline median for the 30–45 min post-second-injection window across saline (n=15, blue), psilocybin (n=21, red), and Ket + Psi (n=8, yellow) experiments.

h, Mean normalized pupil radius (30–45 min post-second injection) per session for each experimental condition; baseline means are shown in gray for all conditions.

i, Mean normalized pupil radius for subjects with paired saline and psilocybin sessions (n=7). Baseline means are shown in gray for both conditions; post-injection means (30–45 min post-second injection) are shown for saline (blue) and psilocybin (red) sessions.

j, Coefficient estimates (error bars: standard error) from a linear mixed-effects model fit to the paired sessions in i, predicting normalized pupil radius from the identity and occurrence of the injection (saline or psilocybin), running speed (z-scored), and time in the experiment (z-scored).

k, Mean normalized pupil radius for subjects with paired saline and Ket + Psi sessions (n=5). Baseline means are shown in gray for both conditions; post-injection means (30–45 min post-second injection) are shown for saline (blue) and Ket + Psi (yellow) sessions.

l, Same as j, but fit to the paired sessions in k, predicting normalized pupil radius from the identity and occurrence of the injection (saline or Ket + Psi), running speed (z-scored), and time in the experiment (z-scored).

Boxplots (f, h, i, k) extend from the first to third quartile with a line at the median; whiskers extend 1.5× the interquartile range beyond the box. Colored asterisks denote significant Wilcoxon signed-rank tests between baseline and post-injection periods within each condition; the black asterisk denotes a significant difference between psilocybin and Ket + Psi conditions (***$p<0.001$, **$p<0.01$, *$p<0.05$). Full statistical details from the LMM are provided in Supplementary Stats_FigS1_Behaviour.xls.

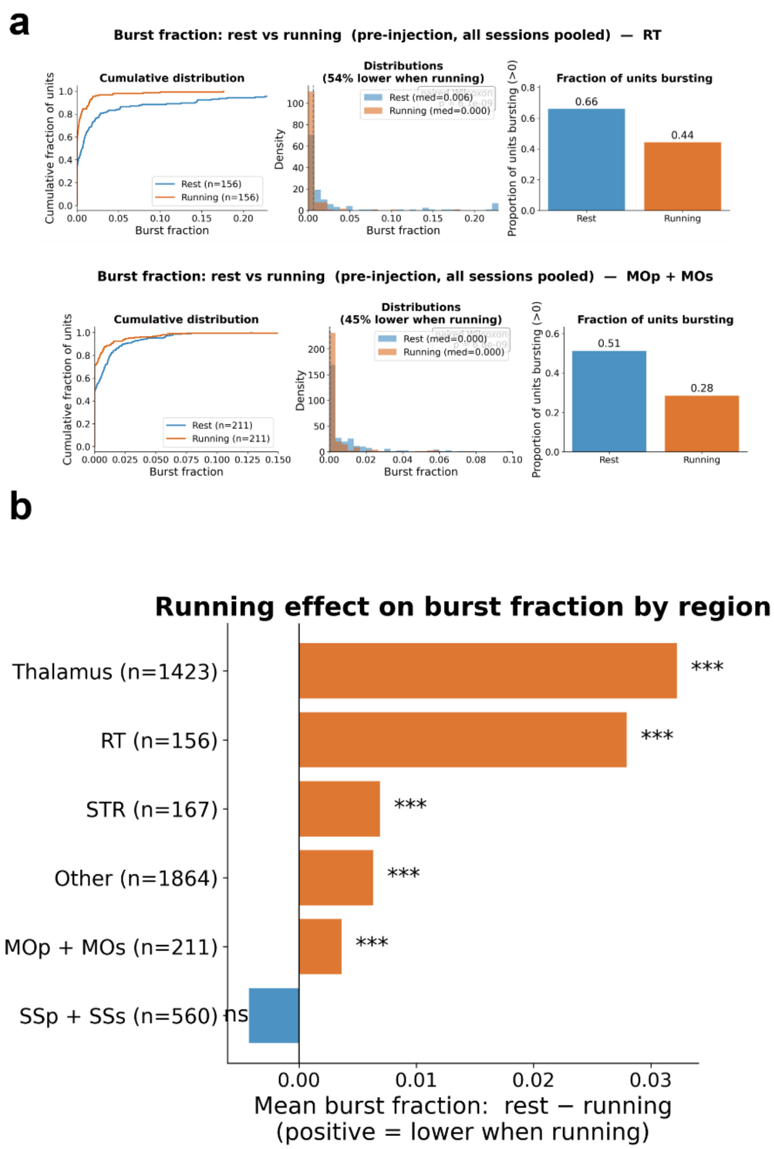


**Figure S2: Burst fraction during rest vs. running, pre-injection-1 window.**

a, Two example regions, RT (top) and MOp + MOs (bottom). Left: cumulative distribution of per-unit burst fraction during rest (blue) and running (orange), n = paired units. Center: histograms of the same data (median values shown in legend); the title gives the percentage of paired units with a lower burst fraction during running than during rest. Right: proportion of units classified as "bursting" (burst_fraction > 0) during rest vs. running.

b, Running effect on burst fraction across regions/groups. Bars show the mean per-unit difference in burst fraction (rest − running); positive = lower burst fraction during running. Asterisks indicate Wilcoxon signed-rank test result (***p < 0.001; ns = not significant); n = number of paired units per region. "Other" = units not assigned to Thalamus, RT, MOp + MOs, SSp + SSs, or STR.

Burst fraction definition and inclusion criteria (all panels): Burst fraction was computed separately for rest and running periods, restricted to the pre- first injection window. A unit receives a usable burst_fraction value for a given state (rest or running) only if it fired ≥ 20 spikes (min_spikes = 20) during that state within this window; otherwise, the value is undefined (NaN). Only units with a valid burst_fraction in both rest and running ("paired units") were included.

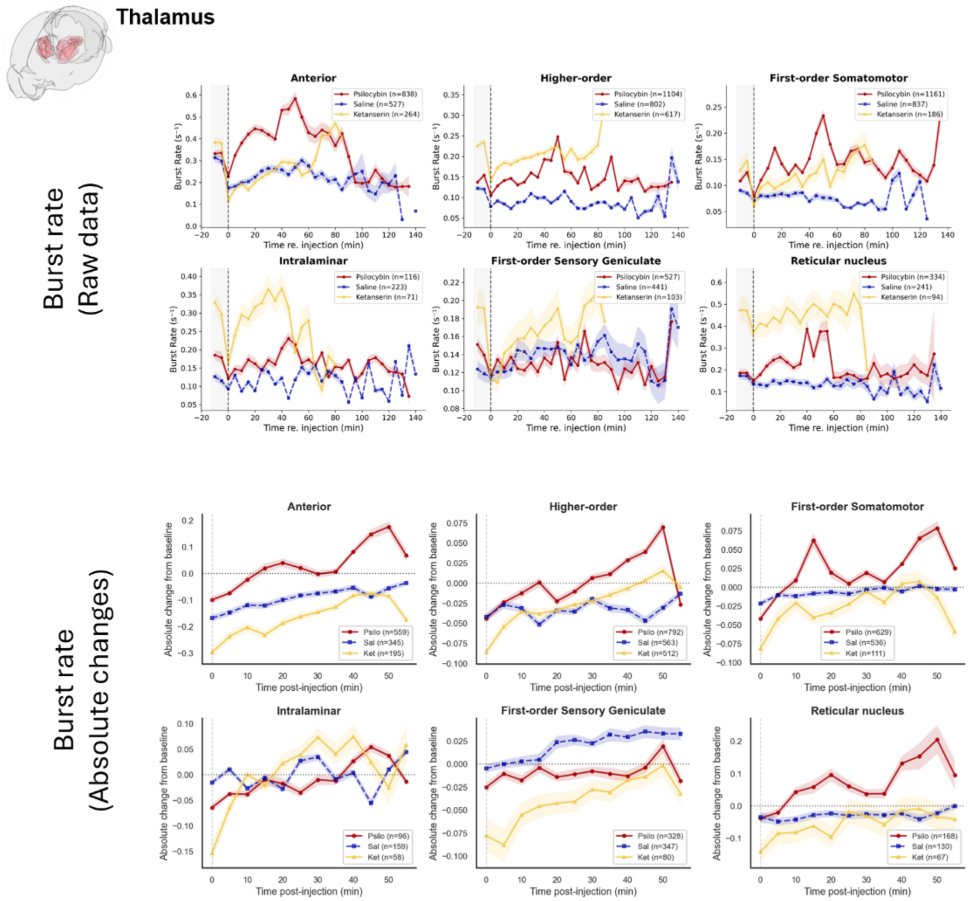


**Figure S3. Burst rate over time in thalamic functional groups (raw data, related to Figure 2c).**

Top: Raw (non-normalized) burst rate, mean ± SEM in 5-min bins, for Psilocybin (red), Saline (blue, dashed), and Ket+Psi (yellow) groups across the full recording session. Gray shading marks the 10-min pre-injection baseline period; the dashed vertical line marks the time of the second injection (t = 0). n indicates the number of units at time 0 per group.

Bottom: Same six groups, showing each unit's burst rate expressed as absolute change from its own baseline (effect − baseline mean), averaged across units (mean ± SEM) in 5-min bins from injection (t = 0) to 60 min post-injection. The dotted horizontal line marks zero change. n is smaller than in the top row because only units with a valid (non-zero) baseline burst fraction are included, as required for per-unit normalization.

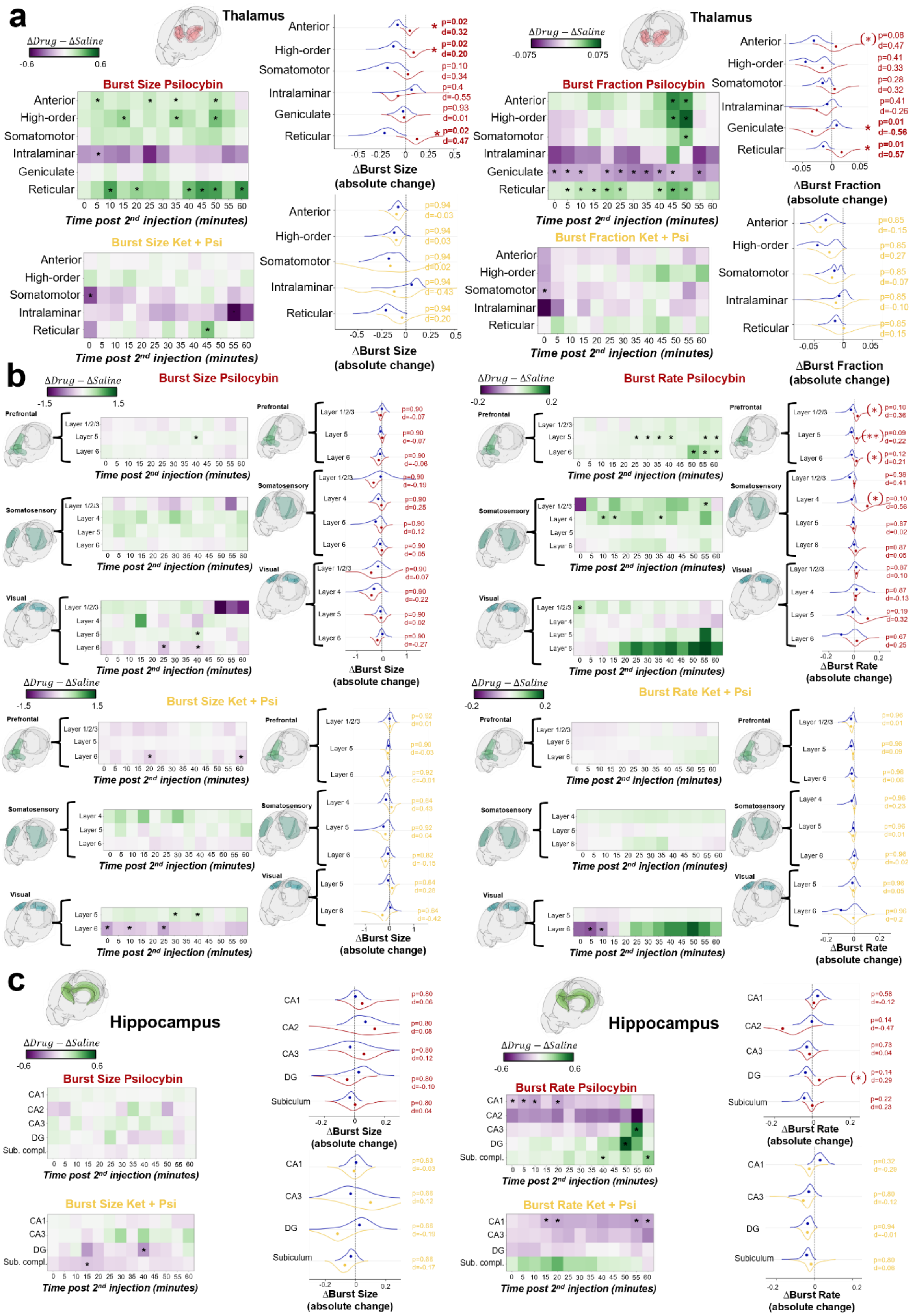

a
Thalamus
ΔDrug − ΔSaline
-0.6
0.6
Burst Size Psilocybin
Anterior
High-order
Somatomotor
Intralaminar
Geniculate
Reticular
0 5 10 15 20 25 30 35 40 45 50 55 60
Time post 2nd injection (minutes)
Burst Size Ket + Psi
p=0.02 d=0.32
p=0.02 d=0.20
p=0.10 d=0.34
p=0.4 d=-0.55
p=0.93 d=0.01
p=0.02 d=0.47
-0.5 -0.25 0 0.25 0.5
ΔBurst Size (absolute change)
p=0.94 d=-0.03
p=0.94 d=0.03
p=0.94 d=0.02
p=0.94 d=-0.43
p=0.94 d=0.20
Thalamus
-0.075
0.075
Burst Fraction Psilocybin
Burst Fraction Ket + Psi
p=0.08 d=0.47
p=0.41 d=0.33
p=0.28 d=0.32
p=0.41 d=-0.26
p=0.01 d=-0.56
p=0.01 d=0.57
-0.05 0 0.05
ΔBurst Fraction (absolute change)
p=0.85 d=-0.15
p=0.85 d=0.27
p=0.85 d=-0.07
p=0.85 d=-0.10
p=0.85 d=0.15
b
-1.5
1.5
Burst Size Psilocybin
Prefrontal
Somatosensory
Visual
Layer 1/2/3
Layer 4
Layer 5
Layer 6
ΔBurst Size (absolute change)
Burst Size Ket + Psi
-0.2
0.2
Burst Rate Psilocybin
Burst Rate Ket + Psi
ΔBurst Rate (absolute change)
c
Hippocampus
Burst Size Psilocybin
Burst Size Ket + Psi
CA1
CA2
CA3
DG
Sub. compl.
Subiculum
Burst Rate Psilocybin
Burst Rate Ket + Psi

**Figure S4. Burst size and burst rate (burst fraction for thalamic groups) dynamics across thalamic groups, cortical layers, and hippocampal subregions.**

For each anatomical grouping — thalamic functional groups (a), isocortical layers across Prefrontal, Somatosensory, and Visual cortex stratified by layer (b), and hippocampal subregions (c) — drug-induced changes in mean burst size and burst rate (burst fraction for thalamic groups) are shown in two complementary formats. Heatmaps show, for each 5-min time bin from 0–60 min post-injection, the difference between groups in the per-unit absolute change from baseline (Δ = effect-period value − baseline value): drug-group Δ minus saline-group Δ. Color follows a diverging purple–green scale centered at zero (purple = relative decrease, green = relative increase under drug); asterisks mark bins where this between-group difference is significant ($p < 0.05$, hierarchical bootstrap, uncorrected for multiple comparisons). Violin plots alongside each heatmap show, for the same rows, the distribution (hierarchical bootstrap, session → unit) of per-unit absolute change pooled over the 40–60 min post-injection window, with dots indicating bootstrap means; p (Benjamini–Hochberg corrected across rows within each panel) and Cohen's d are annotated per row, with the uncorrected p-value shown in parentheses where the effect did not survive FDR correction. For each region grouping, the left pair of panels shows the drug-vs-saline contrast for burst size, and the right for burst rate (burst fraction in a), respectively. Red = Psilocybin, yellow = Ket + Psi, blue = Saline.

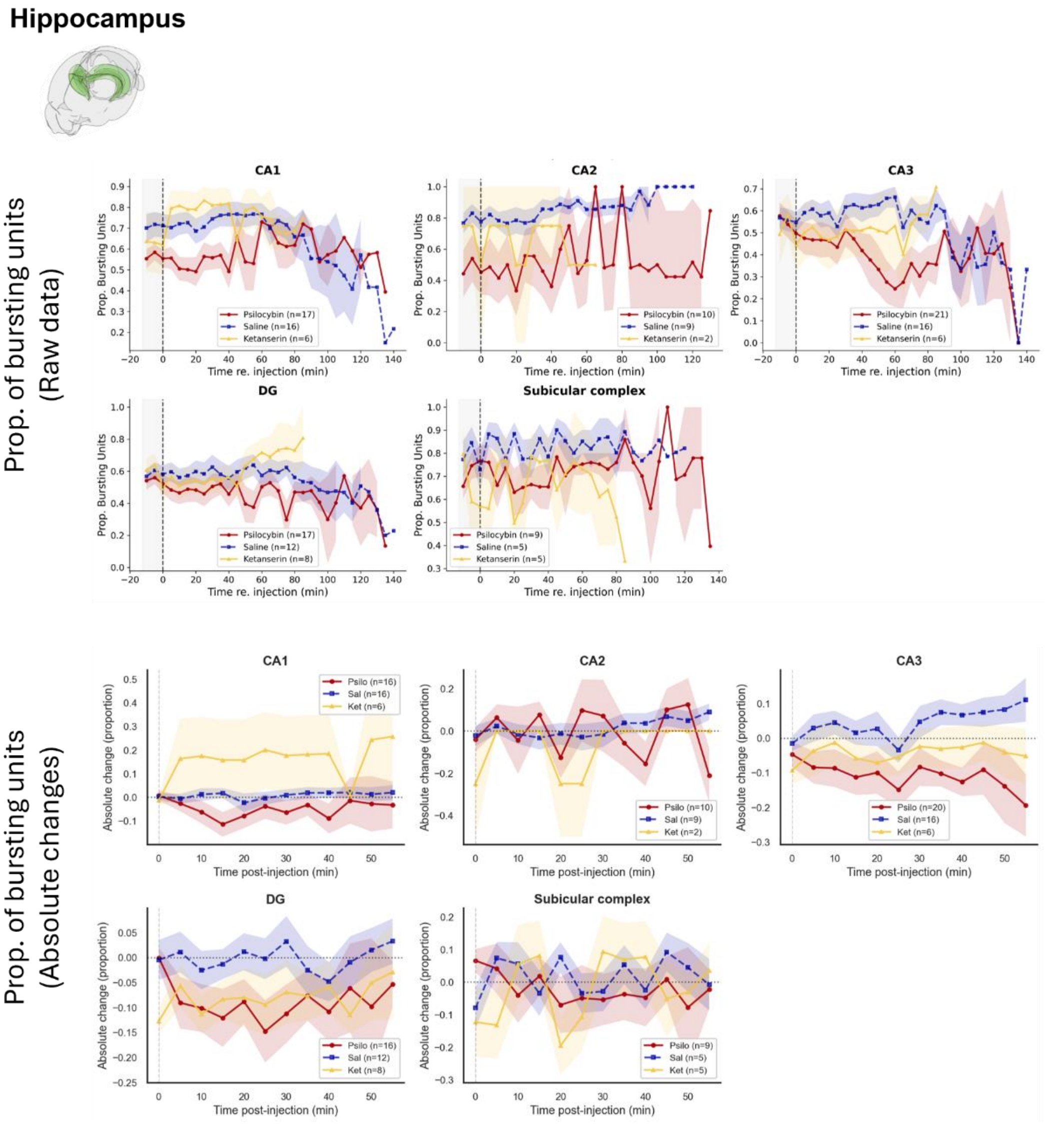


**Figure S5. Raw and baseline-normalized proportion of bursting units in hippocampal areas.** (Top) Raw time courses of the proportion of bursting units in each hippocampal subfield (CA1, CA2, CA3, DG, Subicular complex [SUB + ProS]) before and after injection. The shaded gray region marks the 10-min baseline period; the dashed vertical line marks the time of the second injection (t = 0). Lines show group means ± SEM for Psilocybin (red), Saline (blue, dashed), and Ket + Psi (yellow); n = number of sessions per group, given in each panel's legend. (Bottom) Per-unit absolute change in proportion of bursting units relative to baseline, across the post-injection period (0–60 min, 5-min bins), for the same subfields and groups. Shaded bands = SEM; dotted horizontal line at zero = no change from baseline.

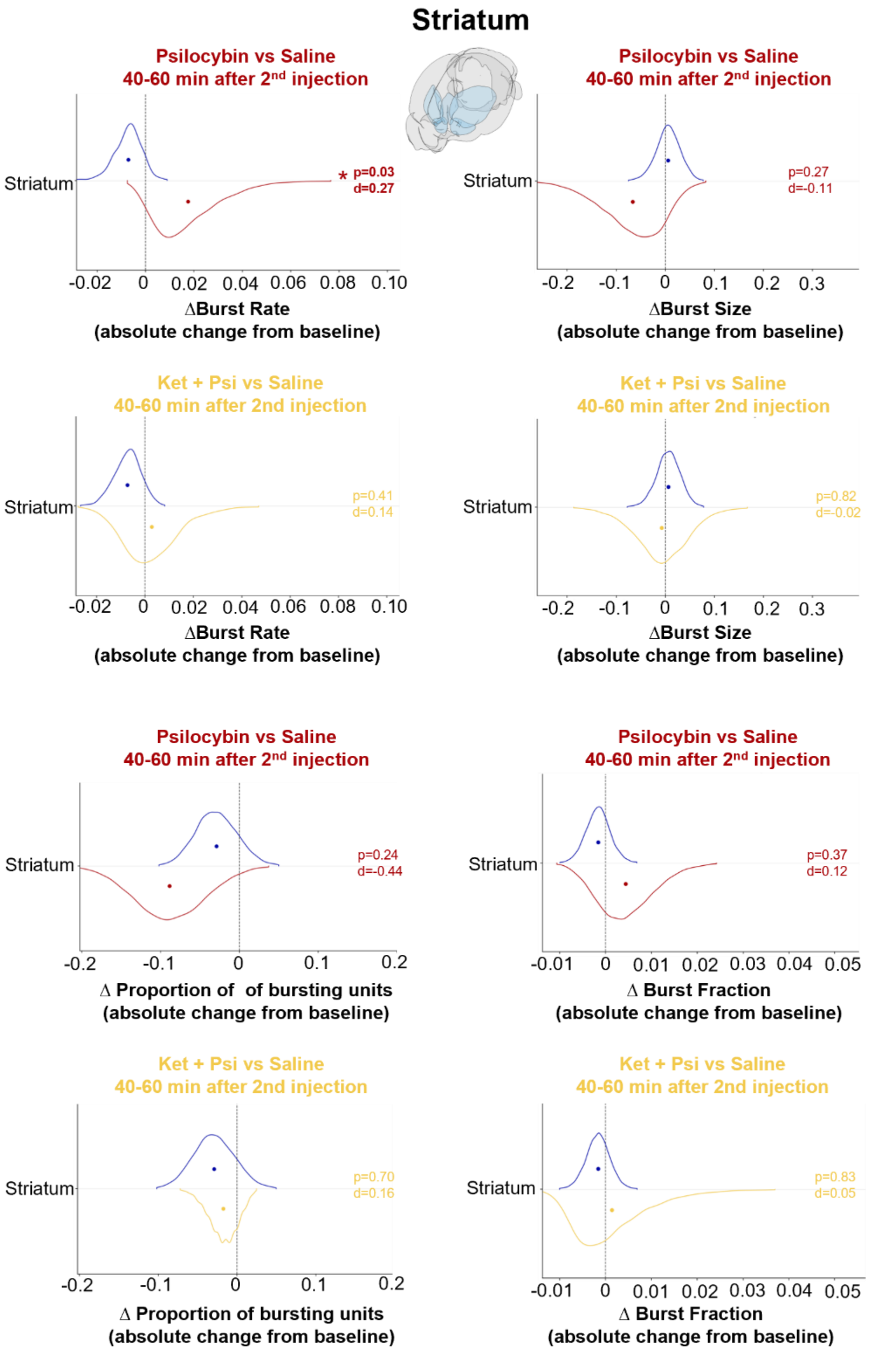


**Figure S6. Burst dynamics in the striatum.** For each of four burst metrics, burst fraction, burst rate ($s^{-1}$), mean burst size (spikes), and proportion of bursting units, violin plots show the distribution (hierarchical bootstrap, session → unit) of the per-unit absolute change from baseline (Δ = 40–60 min post-injection value − baseline value) for the Psilocybin-vs-Saline (top, red vs. blue) and Ket + Psi-vs-Saline (bottom, yellow vs. blue) contrasts. Dots indicate bootstrap means; p and Cohen's d are annotated for the single striatal (STR) row in each panel. Psilocybin significantly increased burst rate (p = 0.03, d = 0.27), an effect abolished by Ketanserin pretreatment (p = 0.41, d = 0.14), consistent with a 5-HT2A-receptor-dependent mechanism. None of the remaining six comparisons reached significance (all p ≥ 0.24).

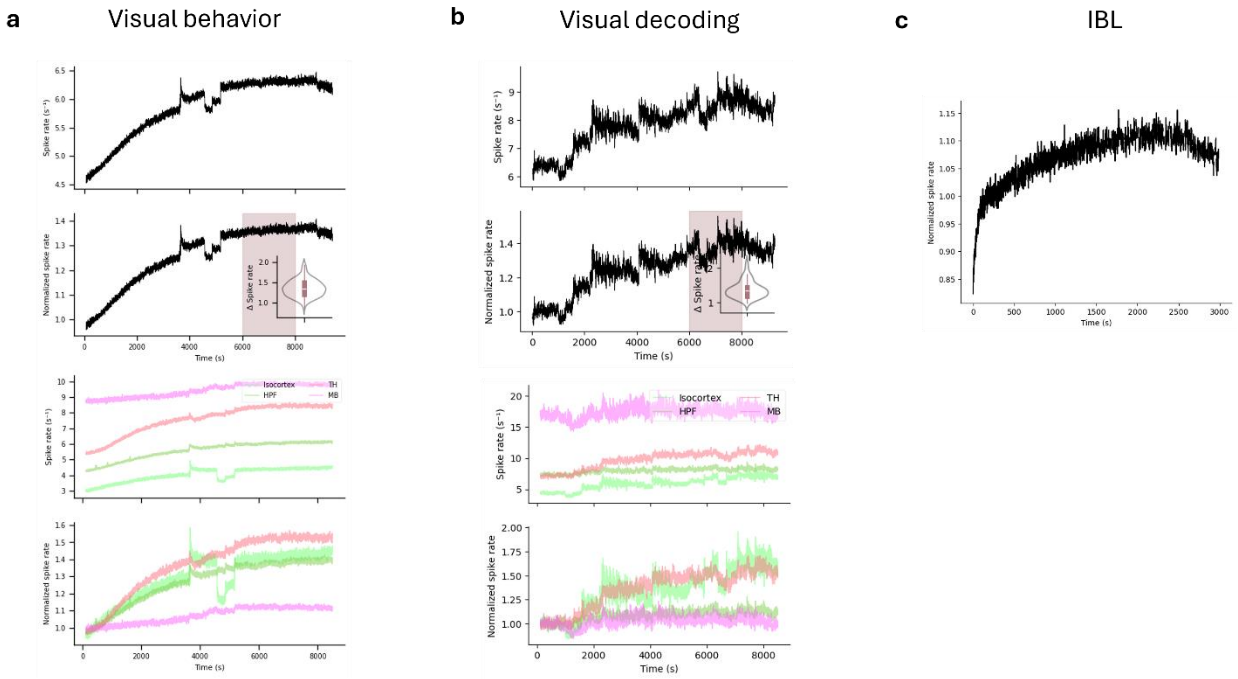


**Figure S7. Firing rate drift in head-fixed Neuropixels recordings.** Gradual increase in population firing rate over the course of head-fixed Neuropixels recording sessions, shown across three independent datasets from publicly available studies in which no injections were delivered.

a, b, Raw (top) and normalized (second row) mean spike rate as a function of recording time for two separate Allen Institute Neuropixels datasets (Siegle et al., 2021). Normalized spike rate increases progressively from session onset, reaching a plateau after approximately 6,000–7,000 s. Inset violin plots show the distribution of Δ spike rate measured in the terminal recording epoch (shaded region) relative to the first 5 min. Lower panels show raw and normalized firing rate broken down by major brain region (Isocortex, Thalamus, HPF, MB), demonstrating that the drift is present across all recorded areas although with different magnitude.

c, A third dataset (International Brain Laboratory., Angelaki, D., Benson, B. *et al.* A brain-wide map of neural activity during complex behaviour. *Nature* **645**, 177–191 (2025)), showing a monotonic increase in normalized firing rate over ~50 min that reaches a stable plateau. Taken together, these data indicate that a gradual upward drift in firing rate is a consistent feature of acute head-fixed Neuropixels recordings even in absence of pharmacological manipulations, likely reflecting cumulative physiological stress responses and/or mechanical settling of the probe, and provides the baseline context for interpreting firing rate time courses in the present study.

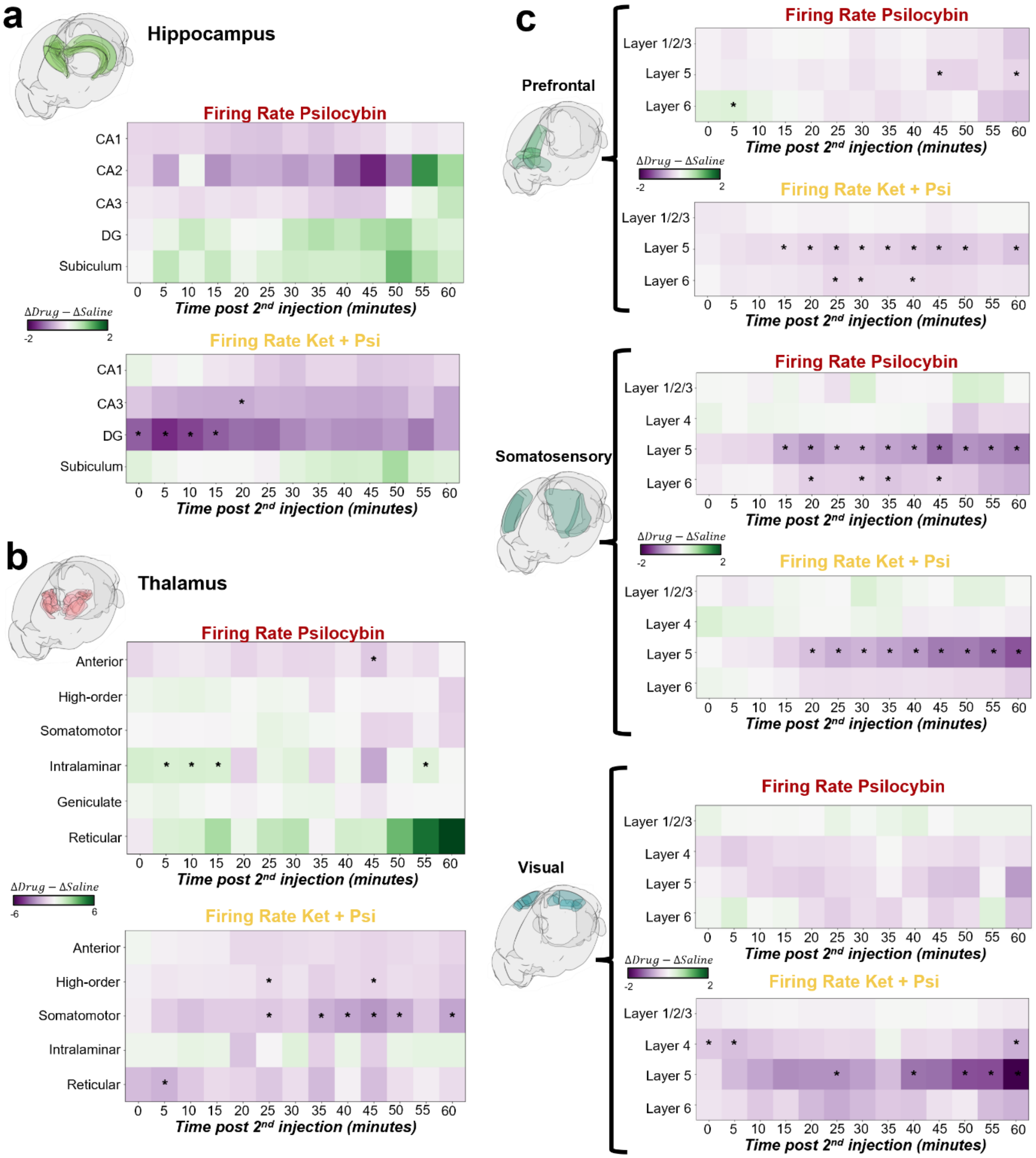


**Figure S8. Drug-induced changes in firing rate (RS units) across brain regions, by time post-injection.** a, Hippocampus (CA1, CA2, CA3, DG, Subiculum). b, Thalamus (Anterior, Higher-order, Somatomotor, Intralaminar, Geniculate, Reticular). c, Isocortex (Prefrontal, Somatosensory, and Visual cortex, stratified by layer). For each anatomical region grouping, heatmaps show, for each 5-min time bin (0–60 min post-injection), the between-group difference in the per-unit absolute change in firing rate from baseline: (Psilocybin or Ket + Psi ΔFR) − (Saline ΔFR), where ΔFR = post-injection firing rate − baseline firing rate (Hz), for RS units only. Color follows a diverging purple–green scale centered at zero: purple = relative decrease in firing rate under drug, green = relative increase. Asterisks mark bins where this between-group difference is significant ($p < 0.05$, hierarchical bootstrap [session → unit], uncorrected for multiple comparisons).

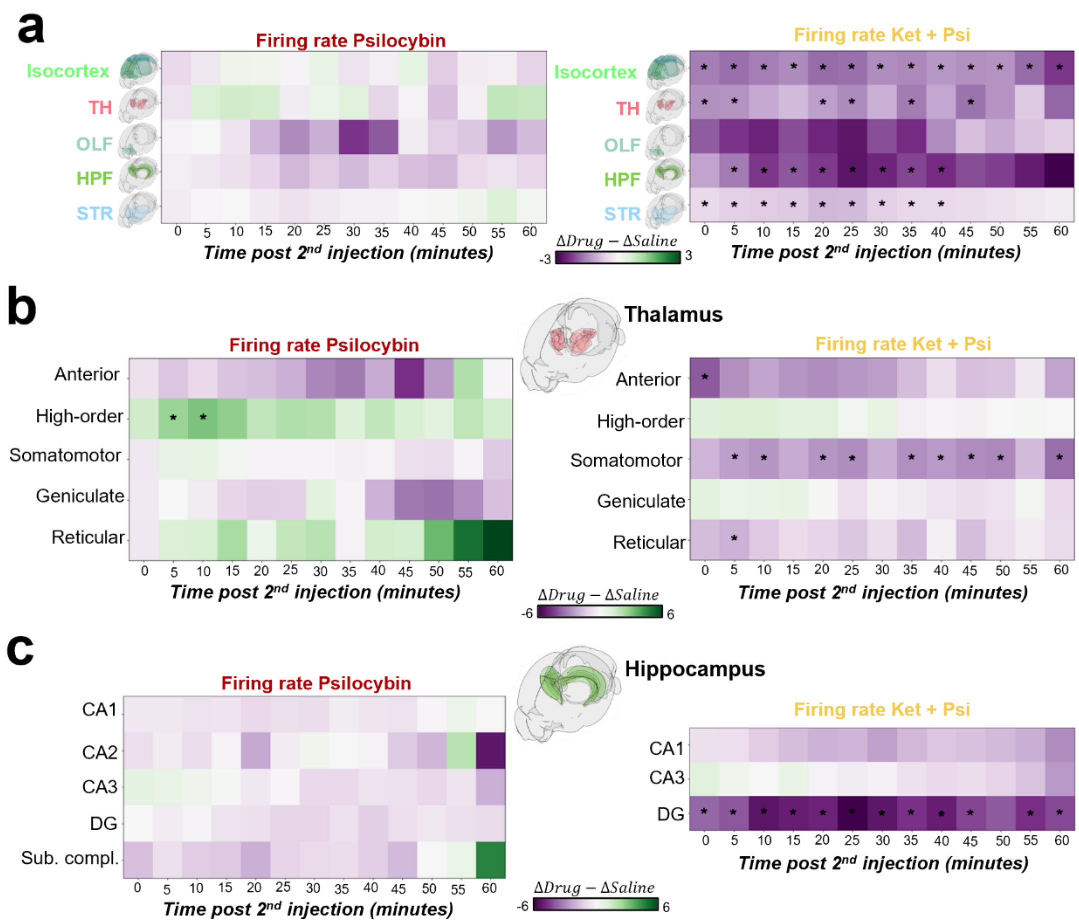


**Figure S9. Drug-induced changes in firing rate (FS units) across brain regions, by time post-injection.** Same representation as in S8 for (a) parent brain regions (Isocortex, Thalamus, OLF, HPF, STR), (b) thalamic nuclei (Anterior, Higher-order, First-order Somatomotor, First-order Sensory Geniculate, Reticular nucleus; Intralaminar excluded due to insufficient FS units), and (c) hippocampal areas.

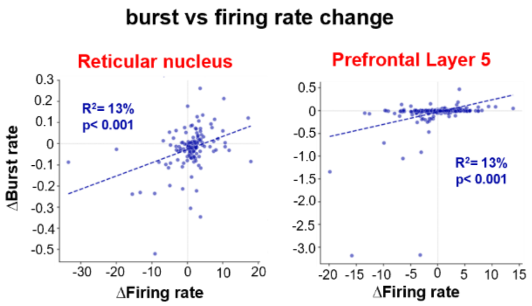

**Figure S10. Correlation analysis in saline group (related to Figure 3e).** Scatter plots of per-unit burst rate change vs. firing rate change in the reticular nucleus (left) and prefrontal cortex layer 5 (right), for the saline condition. $R^2$ and p-values are reported for each region.

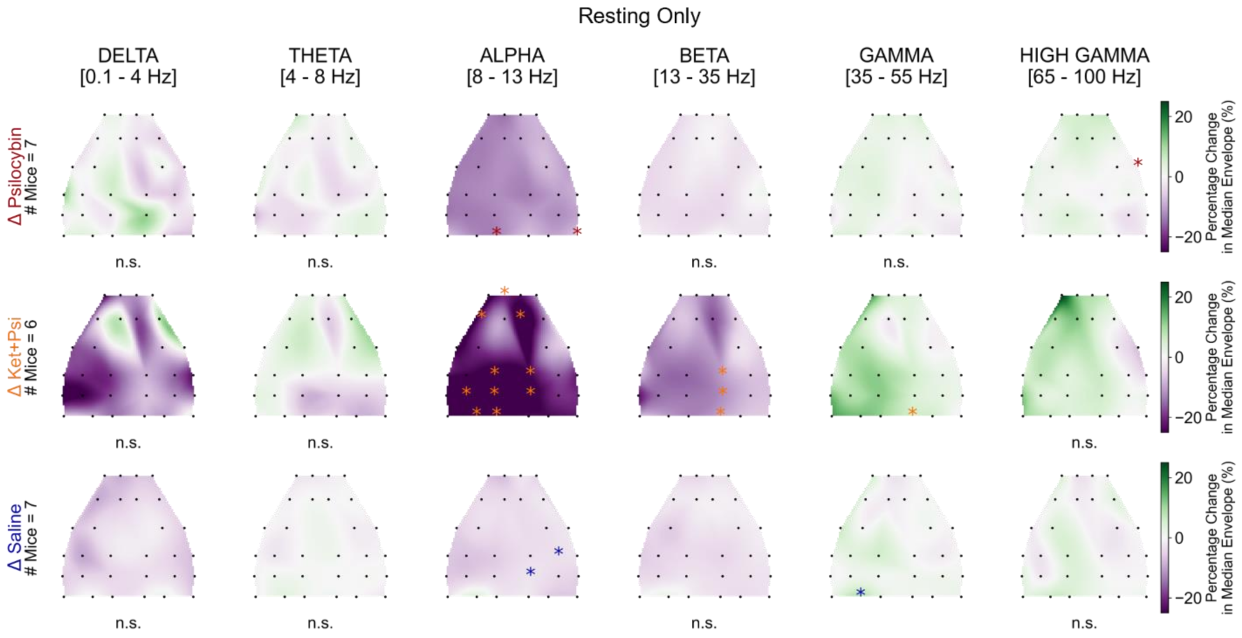


**Figure S11. EEG behavioral control without multiple comparisons.** Topographic plots of the median percentage change in power from baseline under resting state only for psilocybin (top row, N=7), Ket + Psi (middle row, N=6), and saline (bottom row, N=7) conditions, for each frequency band (columns) and EEG channel (black dots). Statistical significance at each channel was assessed using a Wilcoxon ranked-sum test; significant channels are marked with asterisks. Statistical results for each channel are reported in the supplementary file Stats_EEG.xlsx.

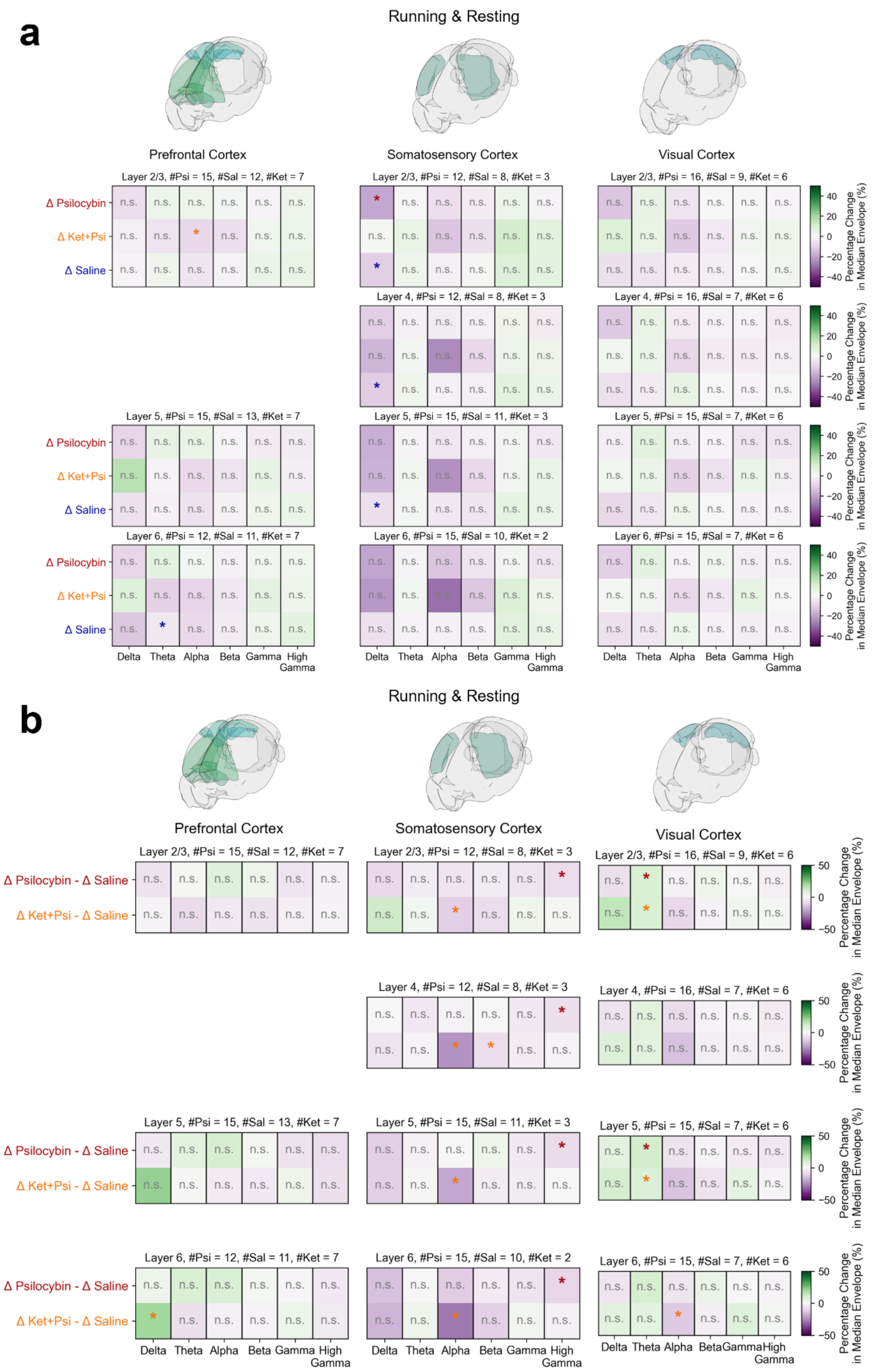

a
Running & Resting
Prefrontal Cortex
Somatosensory Cortex
Visual Cortex
Layer 2/3, #Psi = 15, #Sal = 12, #Ket = 7
Layer 2/3, #Psi = 12, #Sal = 8, #Ket = 3
Layer 2/3, #Psi = 16, #Sal = 9, #Ket = 6
Layer 4, #Psi = 12, #Sal = 8, #Ket = 3
Layer 4, #Psi = 16, #Sal = 7, #Ket = 6
Layer 5, #Psi = 15, #Sal = 13, #Ket = 7
Layer 5, #Psi = 15, #Sal = 11, #Ket = 3
Layer 5, #Psi = 15, #Sal = 7, #Ket = 6
Layer 6, #Psi = 12, #Sal = 11, #Ket = 7
Layer 6, #Psi = 15, #Sal = 10, #Ket = 2
Layer 6, #Psi = 15, #Sal = 7, #Ket = 6
Δ Psilocybin
Δ Ket+Psi
Δ Saline
n.s.
*
Delta
Theta
Alpha
Beta
Gamma
High Gamma
Percentage Change in Median Envelope (%)
40
20
0
−20
−40
b
Running & Resting
Prefrontal Cortex
Somatosensory Cortex
Visual Cortex
Layer 2/3, #Psi = 15, #Sal = 12, #Ket = 7
Layer 2/3, #Psi = 12, #Sal = 8, #Ket = 3
Layer 2/3, #Psi = 16, #Sal = 9, #Ket = 6
Layer 4, #Psi = 12, #Sal = 8, #Ket = 3
Layer 4, #Psi = 16, #Sal = 7, #Ket = 6
Layer 5, #Psi = 15, #Sal = 13, #Ket = 7
Layer 5, #Psi = 15, #Sal = 11, #Ket = 3
Layer 5, #Psi = 15, #Sal = 7, #Ket = 6
Layer 6, #Psi = 12, #Sal = 11, #Ket = 7
Layer 6, #Psi = 15, #Sal = 10, #Ket = 2
Layer 6, #Psi = 15, #Sal = 7, #Ket = 6
Δ Psilocybin - Δ Saline
Δ Ket+Psi - Δ Saline
n.s.
*
Delta
Theta
Alpha
Beta
Gamma
High Gamma
Percentage Change in Median Envelope (%)
50
0
−50

**Figure S12. LFP power analysis confirms the trends observed in the EEG analysis, with somatosensory cortex showing more significant effects than prefrontal and visual areas.**

a, Brain renderings highlighting the cortical regions from which LFPs were recorded (green) for each column: prefrontal cortex, somatosensory cortex, and visual cortex.

b, LFP power analysis across cortical regions (columns: prefrontal, somatosensory, and visual cortex) and cortical layers (rows: layer 2/3, layer 4, layer 5, and layer 6), across all behavioral states (locomotion and rest combined). Each heatmap shows the median percentage change in LFP power from baseline for each frequency band (delta, theta, alpha, beta, gamma, and high gamma), for psilocybin (top row), Ket+ Psi (middle row), and saline (bottom row) conditions. Statistical significance at each frequency band was assessed using a Wilcoxon ranked-sum test; significant bands are marked with asterisks.

c, Scatter plots of median percentage change in LFP power per frequency band (delta to high gamma, left to right) per mouse, averaged across regions and layers, for saline (blue), psilocybin (red), and Ket + Psi (yellow) conditions.

d, Same layout as b, showing the difference in median percentage change in LFP power between drug and saline conditions: psilocybin minus saline (top row) and Ket + Psi minus saline (bottom row).

Statistical significance was assessed using a Mann-Whitney U test; significant bands are marked with asterisks. Detailed statistics are reported in the supplementary file Stats_LFP.xlsx.

**Supplementary Table 1. Unit counts across brain regions and experimental conditions.** Abbreviations: RS, regular-spiking; FS, fast-spiking; OLF, olfactory areas; HPF, hippocampal formation; STR, striatum; RSP, retrosplenial cortex; CA1–CA3, fields of Ammon's horn; DG, dentate gyrus; POST, postsubiculum.

| | Group | Psilocybin | | Saline | | Ket+Psi | | Total | |
|---|---|---|---|---|---|---|---|---|---|
| | | RS | FS | RS | FS | RS | FS | RS | FS |
| Parent Areas | Isocortex | 9,087 | 1,317 | 5,691 | 821 | 2,872 | 463 | 17,650 | 2,601 |
| | Thalamus | 3,977 | 1,187 | 3,017 | 943 | 1,269 | 350 | 8,263 | 2,480 |
| | OLF | 507 | 113 | 862 | 130 | 132 | 16 | 1,501 | 259 |
| | HPF | 2,477 | 615 | 1,303 | 367 | 553 | 111 | 4,333 | 1,093 |
| | STR | 2,235 | | 1,312 | | 666 | | 4,213 | |
| Cortical areas × layer | Prefrontal — Layer 1/2/3 | 987 | 65 | 506 | 37 | 178 | 11 | 1,671 | 113 |
| | Prefrontal — Layer 5 | 2,509 | 231 | 1,806 | 210 | 1,065 | 138 | 5,380 | 579 |
| | Prefrontal — Layer 6 | 1,042 | 152 | 1,173 | 142 | 637 | 103 | 2,852 | 397 |
| | Somatosensory — Layer 1/2/3 | 266 | 47 | 113 | 32 | 37 | 13 | 416 | 92 |
| | Somatosensory — Layer 4 | 419 | 94 | 165 | 46 | 102 | 29 | 686 | 169 |
| | Somatosensory — Layer 5 | 1,039 | 174 | 557 | 101 | 216 | 50 | 1,812 | 325 |
| | Somatosensory — Layer 6 | 1,321 | 309 | 536 | 111 | 235 | 35 | 2,092 | 455 |
| | Visual — Layer 1/2/3 | 172 | 23 | 110 | 16 | 69 | 7 | 351 | 46 |
| | Visual — Layer 4 | 243 | 41 | 90 | 15 | 22 | 8 | 355 | 64 |
| | Visual — Layer 5 | 512 | 77 | 142 | 29 | 99 | 15 | 753 | 121 |
| | Visual — Layer 6 | 414 | 73 | 71 | 20 | 67 | 14 | 552 | 107 |
| | RSP — Layer 1/2/3 | 6 | 1 | 1 | 1 | 7 | 5 | 14 | 7 |
| | RSP — Layer 5 | 21 | 11 | 71 | 6 | 66 | 14 | 158 | 31 |
| | RSP — Layer 6 | 13 | 3 | 65 | 14 | 23 | 7 | 101 | 24 |
| Thalamic groups | Anterior | 838 | 164 | 527 | 73 | 264 | 31 | 1,629 | 268 |
| | Higher-order | 1,104 | 103 | 802 | 53 | 617 | 86 | 2,523 | 242 |
| | First-order Somatomotor | 1,161 | 347 | 837 | 383 | 186 | 89 | 2,184 | 819 |
| | Intralaminar | 116 | 16 | 223 | 2 | 71 | 12 | 410 | 30 |
| | First-order Sensory Geniculate | 527 | 153 | 441 | 153 | 103 | 29 | 1,071 | 335 |
| | Reticular nucleus | 334 | | 241 | | 94 | | 669 | |
| HPF areas | CA1 | 946 | 141 | 389 | 108 | 154 | 22 | 1,489 | 271 |
| | CA3 | 729 | 202 | 397 | 118 | 116 | 42 | 1,242 | 362 |
| | DG | 432 | 199 | 248 | 90 | 151 | 35 | 831 | 324 |
| | Subicular complex | 279 | 48 | 153 | 28 | 128 | 7 | 560 | 83 |
| | CA2 | 55 | 12 | 97 | 18 | 3 | 3 | 155 | 33 |
| | POST | 17 | 9 | 19 | 5 | 1 | 2 | 37 | 16 |

Note: RS/FS classification by waveform duration was not applied to the striatum or reticular nucleus, as this dichotomy assumes co-occurring excitatory and fast-spiking inhibitory populations; striatal neurons are predominantly GABAergic medium spiny neurons, and reticular nucleus neurons are uniformly GABAergic, precluding a meaningful excitatory/inhibitory waveform split in either structure. Unit counts for these two regions are reported as a single merged value spanning the RS and FS columns.

**Supplementary Table 2. Unit counts per 5-min bin for continuous burst metrics (RS units, resting state) for each condition.** Counts reflect unique units per session with rest time and burst fraction > 0 in each bin. Baseline: last 10 min before injection 1; effect period: from injection 2 onward. STR and reticular nucleus include all units regardless of waveform duration as described in Methods.

### Psilocybin

| | Group | Baseline (min before inj 1) | | Effect (min after inj 2) | | | | | | | | | | | |
|---|---|---|---|---|---|---|---|---|---|---|---|---|---|---|---|
| | | *5–10* | *0–5* | *0–5* | *5–10* | *10–15* | *15–20* | *20–25* | *25–30* | *30–35* | *35–40* | *40–45* | *45–50* | *50–55* | *55–60* |
| **Parent areas** | Isocortex | 2,544 | 2,134 | 1,675 | 1,445 | 1,451 | 1,334 | 1,390 | 1,425 | 1,349 | 1,381 | 1,361 | 680 | 587 | 536 |
| | Thalamus | 3,019 | 2,811 | 2,729 | 2,450 | 2,466 | 2,300 | 2,466 | 2,379 | 2,408 | 2,385 | 2,291 | 1,568 | 1,278 | 1,214 |
| | OLF | 228 | 164 | 161 | 128 | 106 | 103 | 109 | 109 | 118 | 104 | 112 | 71 | 69 | 61 |
| | HPF | 1,498 | 1,409 | 1,354 | 1,112 | 1,099 | 1,038 | 1,039 | 1,037 | 999 | 1,005 | 933 | 461 | 344 | 321 |
| | STR | 849 | 760 | 545 | 460 | 446 | 448 | 451 | 414 | 405 | 421 | 410 | 174 | 135 | 133 |
| **Cortical Areas × Layers** | Prefrontal — Layer 1/2/3 | 128 | 90 | 73 | 59 | 59 | 50 | 60 | 63 | 62 | 50 | 47 | 34 | 24 | 23 |
| | Prefrontal — Layer 5 | 742 | 645 | 517 | 471 | 439 | 390 | 426 | 466 | 440 | 463 | 425 | 237 | 208 | 189 |
| | Prefrontal — Layer 6 | 389 | 349 | 284 | 233 | 248 | 243 | 240 | 216 | 204 | 202 | 234 | 96 | 75 | 64 |
| | Somatosensory — Layer 1/2/3 | 36 | 27 | 14 | 15 | 13 | 8 | 12 | 11 | 11 | 7 | 8 | 4 | 5 | 4 |
| | Somatosensory — Layer 4 | 84 | 62 | 45 | 30 | 36 | 20 | 29 | 35 | 25 | 29 | 37 | 29 | 24 | 25 |
| | Somatosensory — Layer 5 | 343 | 286 | 221 | 182 | 199 | 188 | 184 | 189 | 183 | 194 | 201 | 92 | 86 | 79 |
| | Somatosensory — Layer 6 | 462 | 392 | 295 | 264 | 255 | 239 | 244 | 248 | 237 | 255 | 228 | 80 | 78 | 69 |
| | Visual — Layer 1/2/3 | 23 | 20 | 19 | 16 | 19 | 22 | 16 | 17 | 15 | 14 | 10 | 4 | 3 | 4 |
| | Visual — Layer 4 | 32 | 13 | 15 | 17 | 13 | 14 | 14 | 16 | 18 | 14 | 15 | 7 | 7 | 6 |
| | Visual — Layer 5 | 122 | 99 | 81 | 67 | 71 | 62 | 67 | 73 | 69 | 66 | 70 | 48 | 36 | 32 |
| | Visual — Layer 6 | 141 | 114 | 97 | 71 | 76 | 78 | 79 | 75 | 70 | 68 | 67 | 32 | 26 | 25 |
| | RSP — Layer 1/2/3 | 4 | 2 | | 1 | 4 | 2 | 1 | 2 | 2 | 3 | 1 | | | 1 |
| | RSP — Layer 5 | 7 | 7 | 5 | 6 | 6 | 3 | 5 | 5 | 5 | 5 | 2 | 3 | 2 | 3 |
| | RSP — Layer 6 | 1 | 1 | 1 | 2 | 2 | 1 | 1 | 1 | 1 | 1 | | | | |
| **Thalamic groups** | Anterior | 597 | 569 | 559 | 428 | 425 | 394 | 418 | 403 | 405 | 395 | 385 | 253 | 196 | 188 |
| | Higher-order | 876 | 790 | 792 | 716 | 722 | 657 | 740 | 690 | 714 | 713 | 655 | 448 | 321 | 297 |
| | First-order Somatomotor | 725 | 702 | 629 | 595 | 587 | 606 | 578 | 579 | 592 | 605 | 612 | 328 | 271 | 273 |
| | Intralaminar | 99 | 99 | 96 | 97 | 98 | 95 | 99 | 98 | 92 | 96 | 97 | 94 | 94 | 96 |
| | First-order Sensory Geniculate | 362 | 317 | 328 | 314 | 322 | 309 | 311 | 303 | 295 | 291 | 276 | 243 | 210 | 189 |
| | Reticular nucleus | 187 | 177 | 168 | 150 | 163 | 142 | 175 | 156 | 164 | 138 | 127 | 77 | 62 | 49 |

| | | | | | | | | | | | | | | | |
|---|---|---|---|---|---|---|---|---|---|---|---|---|---|---|---|
| HPF areas | CA1 | 541 | 514 | 484 | 399 | 371 | 351 | 350 | 374 | 349 | 385 | 332 | 212 | 143 | 146 |
| | CA3 | 458 | 425 | 410 | 326 | 333 | 309 | 300 | 289 | 271 | 263 | 288 | 70 | 60 | 57 |
| | DG | 242 | 241 | 222 | 187 | 184 | 174 | 186 | 178 | 178 | 168 | 157 | 68 | 57 | 49 |
| | Subicular complex | 204 | 176 | 196 | 170 | 174 | 170 | 169 | 159 | 164 | 148 | 132 | 94 | 67 | 50 |
| | CA2 | 38 | 37 | 36 | 25 | 22 | 25 | 23 | 26 | 24 | 25 | 24 | 17 | 17 | 18 |
| | POST | 9 | 12 | 6 | 3 | 10 | 6 | 9 | 8 | 10 | 11 | | | | |

### Saline

| | Group | Baseline (min before inj 1) | | Effect (min after inj 2) | | | | | | | | | | | |
|---|---|---|---|---|---|---|---|---|---|---|---|---|---|---|---|
| | | ***5–10*** | ***0–5*** | ***0–5*** | ***5–10*** | ***10–15*** | ***15–20*** | ***20–25*** | ***25–30*** | ***30–35*** | ***35–40*** | ***40–45*** | ***45–50*** | ***50–55*** | ***55–60*** |
| Parent areas | Isocortex | 1,874 | 1,831 | 1,235 | 1,322 | 1,367 | 1,344 | 1,314 | 1,275 | 1,316 | 1,182 | 1,080 | 1,093 | 1,064 | 1,029 |
| | Thalamus | 2,557 | 2,477 | 2,210 | 2,251 | 2,264 | 2,284 | 2,245 | 2,281 | 2,218 | 2,084 | 2,003 | 1,959 | 2,003 | 2,006 |
| | OLF | 374 | 350 | 268 | 271 | 289 | 295 | 286 | 291 | 306 | 276 | 265 | 273 | 292 | 287 |
| | HPF | 854 | 830 | 756 | 763 | 759 | 758 | 757 | 739 | 765 | 715 | 686 | 685 | 682 | 678 |
| | STR | 557 | 531 | 392 | 391 | 426 | 407 | 386 | 405 | 360 | 297 | 269 | 254 | 272 | 281 |
| Cortical Areas × Layers | Prefrontal — Layer 1/2/3 | 109 | 118 | 68 | 82 | 77 | 82 | 76 | 79 | 65 | 58 | 60 | 56 | 63 | 53 |
| | Prefrontal — Layer 5 | 679 | 659 | 460 | 491 | 505 | 506 | 498 | 492 | 503 | 444 | 418 | 417 | 394 | 378 |
| | Prefrontal — Layer 6 | 379 | 360 | 256 | 272 | 274 | 266 | 261 | 251 | 244 | 238 | 194 | 215 | 192 | 191 |
| | Somatosensory — Layer 1/2/3 | 24 | 23 | 19 | 17 | 14 | 14 | 15 | 15 | 15 | 11 | 10 | 15 | 15 | 16 |
| | Somatosensory — Layer 4 | 47 | 37 | 25 | 28 | 26 | 27 | 23 | 21 | 30 | 32 | 28 | 26 | 26 | 26 |
| | Somatosensory — Layer 5 | 197 | 202 | 118 | 121 | 125 | 125 | 111 | 111 | 123 | 111 | 103 | 100 | 101 | 103 |
| | Somatosensory — Layer 6 | 185 | 171 | 118 | 114 | 133 | 125 | 121 | 105 | 124 | 86 | 87 | 97 | 96 | 96 |
| | Visual — Layer 1/2/3 | 9 | 11 | 8 | 10 | 8 | 6 | 10 | 8 | 10 | 10 | 8 | 9 | 9 | 8 |
| | Visual — Layer 4 | 23 | 20 | 14 | 17 | 19 | 14 | 16 | 16 | 18 | 18 | 20 | 20 | 19 | 19 |
| | Visual — Layer 5 | 60 | 66 | 41 | 48 | 52 | 55 | 56 | 49 | 54 | 51 | 57 | 51 | 55 | 52 |
| | Visual — Layer 6 | 30 | 32 | 19 | 22 | 24 | 25 | 25 | 22 | 27 | 24 | 25 | 26 | 27 | 26 |
| | RSP — Layer 1/2/3 | 1 | 1 | 1 | 1 | 1 | 1 | 1 | 1 | 1 | 1 | 1 | 1 | 1 | 1 |
| | RSP — Layer 5 | 11 | 15 | 5 | 9 | 9 | 8 | 8 | 9 | 12 | 9 | 8 | 5 | 6 | 8 |
| | RSP — Layer 6 | 17 | 12 | 11 | 8 | 10 | 8 | 9 | 9 | 12 | 14 | 10 | 10 | 7 | 8 |
| Thalamic groups | Anterior | 413 | 406 | 345 | 371 | 375 | 377 | 373 | 366 | 361 | 337 | 321 | 313 | 313 | 306 |
| | Higher-order | 665 | 631 | 563 | 595 | 593 | 584 | 580 | 595 | 478 | 434 | 406 | 398 | 413 | 411 |
| | First-order Somatomotor | 645 | 639 | 536 | 537 | 532 | 555 | 534 | 558 | 577 | 534 | 517 | 493 | 497 | 506 |
| | Intralaminar | 163 | 145 | 159 | 145 | 156 | 150 | 152 | 140 | 158 | 143 | 141 | 131 | 161 | 160 |

| | | | | | | | | | | | | | | | |
|---|---|---|---|---|---|---|---|---|---|---|---|---|---|---|---|
| | First-order Sensory Geniculate | 364 | 344 | 347 | 337 | 350 | 347 | 333 | 342 | 354 | 352 | 343 | 352 | 338 | 345 |
| | Reticular nucleus | 139 | 145 | 130 | 126 | 134 | 128 | 137 | 125 | 131 | 127 | 115 | 114 | 118 | 119 |
| **HPF areas** | CA1 | 264 | 257 | 245 | 242 | 245 | 240 | 235 | 236 | 246 | 234 | 229 | 230 | 226 | 228 |
| | CA3 | 242 | 242 | 208 | 209 | 212 | 208 | 209 | 201 | 205 | 189 | 165 | 155 | 159 | 155 |
| | DG | 139 | 123 | 120 | 113 | 112 | 116 | 114 | 107 | 118 | 96 | 94 | 101 | 109 | 106 |
| | Subicular complex | 121 | 122 | 104 | 117 | 109 | 110 | 117 | 113 | 114 | 112 | 117 | 115 | 105 | 104 |
| | CA2 | 71 | 67 | 63 | 65 | 64 | 65 | 65 | 65 | 65 | 66 | 65 | 67 | 65 | 68 |
| | POST | 17 | 19 | 16 | 17 | 17 | 19 | 17 | 17 | 17 | 18 | 16 | 17 | 18 | 17 |

### Ket + Psi

| | **Group** | **Baseline (min before inj 1)** | | **Effect (min after inj 2)** | | | | | | | | | | | |
|---|---|---|---|---|---|---|---|---|---|---|---|---|---|---|---|
| | | ***5–10*** | ***0–5*** | ***0–5*** | ***5–10*** | ***10–15*** | ***15–20*** | ***20–25*** | ***25–30*** | ***30–35*** | ***35–40*** | ***40–45*** | ***45–50*** | ***50–55*** | ***55–60*** |
| **Parent areas** | Isocortex | 1,012 | 988 | 719 | 772 | 792 | 799 | 821 | 853 | 887 | 833 | 850 | 862 | 508 | 479 |
| | Thalamus | 1,137 | 1,082 | 1,049 | 1,054 | 1,098 | 1,091 | 1,079 | 1,086 | 1,101 | 1,112 | 1,127 | 1,138 | 856 | 730 |
| | OLF | 62 | 57 | 42 | 40 | 41 | 41 | 57 | 47 | 49 | 46 | 49 | 48 | 38 | 37 |
| | HPF | 369 | 367 | 322 | 330 | 324 | 327 | 320 | 334 | 331 | 331 | 335 | 332 | 246 | 218 |
| | STR | 259 | 241 | 177 | 188 | 200 | 190 | 196 | 194 | 200 | 197 | 215 | 210 | 115 | 94 |
| **Cortical Areas × Layers** | Prefrontal — Layer 1/2/3 | 55 | 64 | 33 | 37 | 36 | 33 | 41 | 42 | 47 | 45 | 41 | 44 | 38 | 34 |
| | Prefrontal — Layer 5 | 428 | 409 | 309 | 327 | 315 | 331 | 341 | 366 | 371 | 346 | 346 | 355 | 180 | 169 |
| | Prefrontal — Layer 6 | 210 | 189 | 137 | 143 | 140 | 146 | 152 | 153 | 165 | 155 | 164 | 168 | 94 | 93 |
| | Somatosensory — Layer 1/2/3 | 4 | 8 | 5 | 4 | 4 | 4 | 4 | 4 | 4 | 7 | 5 | 4 | 6 | 4 |
| | Somatosensory — Layer 4 | 26 | 27 | 27 | 28 | 30 | 26 | 31 | 28 | 26 | 24 | 25 | 23 | 18 | 14 |
| | Somatosensory — Layer 5 | 76 | 81 | 58 | 69 | 73 | 73 | 74 | 70 | 81 | 74 | 76 | 72 | 51 | 52 |
| | Somatosensory — Layer 6 | 110 | 103 | 80 | 91 | 103 | 92 | 101 | 101 | 100 | 95 | 103 | 103 | 63 | 57 |
| | Visual — Layer 1/2/3 | 1 | 5 | 3 | 2 | 2 | 3 | 4 | 4 | 3 | 4 | 2 | 3 | 3 | 5 |
| | Visual — Layer 4 | 4 | 3 | 2 | 3 | 2 | 2 | 2 | 2 | 3 | 3 | 4 | 3 | 1 | |
| | Visual — Layer 5 | 25 | 30 | 20 | 18 | 22 | 21 | 20 | 23 | 24 | 20 | 23 | 24 | 13 | 17 |
| | Visual — Layer 6 | 28 | 26 | 18 | 18 | 25 | 26 | 20 | 26 | 27 | 26 | 28 | 27 | 21 | 14 |
| | RSP — Layer 1/2/3 | 2 | 1 | | 1 | 2 | 1 | | | | | 1 | 1 | 1 | |
| | RSP — Layer 5 | 19 | 20 | 12 | 13 | 17 | 20 | 14 | 17 | 17 | 16 | 15 | 16 | 2 | 2 |

| | | | | | | | | | | | | | | | |
|---|---|---|---|---|---|---|---|---|---|---|---|---|---|---|---|
| | RSP — Layer 6 | 7 | 6 | 4 | 5 | 5 | 6 | 4 | 3 | 5 | 3 | 2 | 2 | 2 | 2 |
| Thalamic groups | Anterior | 225 | 229 | 195 | 204 | 212 | 209 | 219 | 219 | 220 | 219 | 220 | 229 | 188 | 148 |
| | Higher-order | 542 | 501 | 512 | 512 | 535 | 536 | 519 | 526 | 535 | 530 | 546 | 548 | 391 | 302 |
| | First-order Somatomotor | 129 | 111 | 111 | 106 | 114 | 108 | 105 | 103 | 108 | 124 | 120 | 120 | 81 | 86 |
| | Intralaminar | 60 | 59 | 58 | 60 | 60 | 60 | 60 | 60 | 60 | 60 | 60 | 60 | 17 | 16 |
| | First-order Sensory Geniculate | 81 | 83 | 80 | 77 | 79 | 82 | 81 | 82 | 83 | 82 | 84 | 84 | 84 | 84 |
| | Reticular nucleus | 73 | 73 | 67 | 68 | 71 | 69 | 68 | 70 | 69 | 71 | 71 | 70 | 69 | 67 |
| HPF areas | CA1 | 101 | 102 | 91 | 94 | 98 | 95 | 98 | 102 | 97 | 97 | 97 | 95 | 71 | 73 |
| | CA3 | 80 | 78 | 71 | 72 | 70 | 69 | 68 | 69 | 71 | 70 | 73 | 71 | 35 | 35 |
| | DG | 87 | 82 | 60 | 67 | 61 | 63 | 64 | 65 | 66 | 64 | 66 | 60 | 47 | 41 |
| | Subicular complex | 98 | 103 | 99 | 95 | 93 | 98 | 89 | 96 | 95 | 98 | 97 | 104 | 92 | 68 |
| | CA2 | 2 | 2 | 1 | 2 | 2 | 2 | 1 | 1 | 2 | 2 | 2 | 2 | 1 | 1 |
| | POST | 1 | | | | | | | 1 | | | | | | |
